# Biosynthetic Olfactory System from Multiplexed Engineered Microbes (BOSMEM)

**DOI:** 10.64898/2026.09.25.753839

**Authors:** Hann Tu, Shohreh Vanaei, Balasubalakshmi Balasubramanian, Pranav Mundada, Rucha Deshpande, Swaralee Kulkarni, Itishree Sahoo, Neel Joshi

**Affiliations:** Department of Bioengineering, Northeastern University; Department of Chemistry and Chemical Biology, Northeastern University; Department of Chemistry, Tufts University; Department of Biomedical Engineering, Tufts University

## Abstract

Whole-cell biosensors can detect many types of chemicals in complex environments, but existing designs force a tradeoff in multiplexed sensing: separating sensor strains into individual reservoirs limits how many analytes can be monitored at once, while pooling strains in open co-culture lets faster-growing strains take over within days, corrupting the signal. The microfluidic “mother machine” has been used for over a decade, mostly to study individual bacterial strains with single-cell resolution, but never to house multiple engineered strains simultaneously for sensing applications. We repurpose this device to hold a library of genetically distinct biosensor strains, physically isolating each strain in its own growth channel while continuous flow keeps every strain fed. Each strain reports on a target chemical through a unique combination of three fluorescent proteins, so seven distinguishable signals emerge from only three optical channels. We characterized turn-on and turn-off kinetics of several biosensors in the mother machine. Over a multi-day experiment with repeated chemical exposures, biosensors housed this way matched the expected on/off state and held stable population ratios, while biosensors in an open batch co-culture drifted and lost reliable signal. This establishes mother-machine-based multiplexing as a scalable route to long-term chemical monitoring with living sensors.

## Introduction

The accelerating pace of global human impact has intensified the need for continuous, multiplexed chemical monitoring across environmental, food safety and public health settings^1^. Current detection technologies face fundamental constraints preventing real-time, field-deployable operation^2^. While advanced analytical instruments offer high specificity, they require time-consuming sample preparation and laboratory infrastructure^3^. Furthermore, protein and nucleic acid-based biorecognition^4,5^ elements degrade within hours to days under continuous operation, and abiotic chemical sensors suffer from signal drift^6^ and cross-reactivity^7^, restricting reliable performance in heterogeneous matrices. Olfactory-mimetic platforms^8^, such as chemical sensor arrays, incorporate multiple receptors or sensing elements to achieve pattern-recognition detection that can distinguish structurally similar analytes. However, these platforms^9^ still lack the multiplexing^10^ and real-time detection required for complex samples, failing to replicate the discernment, continuity, and portability of true olfaction^11^. Synthetic biology can partially address these gaps by engineering microbes as whole-cell biosensors^12^ (WCBs) capable of detecting chemicals in complex environments via analyte-inducible genetic circuits that produce optical, acoustic, or electrical output signals^13,14^.

Despite this promise, broader WCB deployment lags due to three persistent limitations. While WCBs have been integrated into paper-based devices^15^, hydrogels^16^, and microfluidic chips^17^, spatial separation of biosensor populations into discrete compartments limits multiplexing capability, as each new analyte channel requires a physically distinct reservoir^18^. Additionally, such platforms require specialized storage where cells experience a period of dormancy before activation, which in turn hinders fast response times. Conversely, batch co-culture formats allow unrestricted inter-strain resource competition^19^, driving population shifts that erode sensing reliability over time. Consequently, no existing platform has demonstrated simultaneous, reversible turn-on and turn-off detection across a library of more than a few biosensors operating continuously over multiple days.

The mother machine^20^ is a microfluidic device in which bacteria grow in thousands of narrow dead-end channels fed by a single central media channel, enabling steady continuous cell growth under constant nutrient flow over many days. Each growth channel physically isolates a resident “mother cell” from the others, eliminating inter-cell resource competition that can destabilize batch co-cultures. The mother machine has been used mostly to study fundamental aspects of cell biology,^20,21^ including cell-size fluctuations, cellular aging, and gene expression dynamics^22^. The properties of the mother machine that are useful for microbiology studies – enabling single-cell tracking, minimizing competition between cells, relatively inexpensive fabrication – can also be adapted to biotechnological applications. In this work, we seek to repurpose the mother machine as a platform for highly multiplexed WCB deployment.

Here we present the Biosynthetic Olfactory System from Multiplexed Engineered Microbes (BOSMEM), a detection platform that deploys a library of genetically distinct WCBs within a single mother machine device (Fig. 1). Each biosensor produces a unique spectral code through combinatorial expression of multiple fluorescent proteins, exponentially expanding the number of distinguishable activation signatures from a small number of optical channels. The physical partitioning of growth channels eliminates inter-strain resource competition and sustains all biosensor strains at peak metabolic activity under continuous nutrient flow, enabling a large sensing population to operate within a compact footprint over many days. This architecture creates a structural basis for pattern-recognition-based analyte identification across the sensor ensemble, conceptually analogous to the combinatorial strategy of the mammalian olfactory system^23^. We demonstrate reversible turn-on and turn-off detection kinetics, stable population distributions and sustained biosensor functionality over the course of almost 2 weeks, representing a new biotechnological application of the mother machine.

**Fig. 1.**
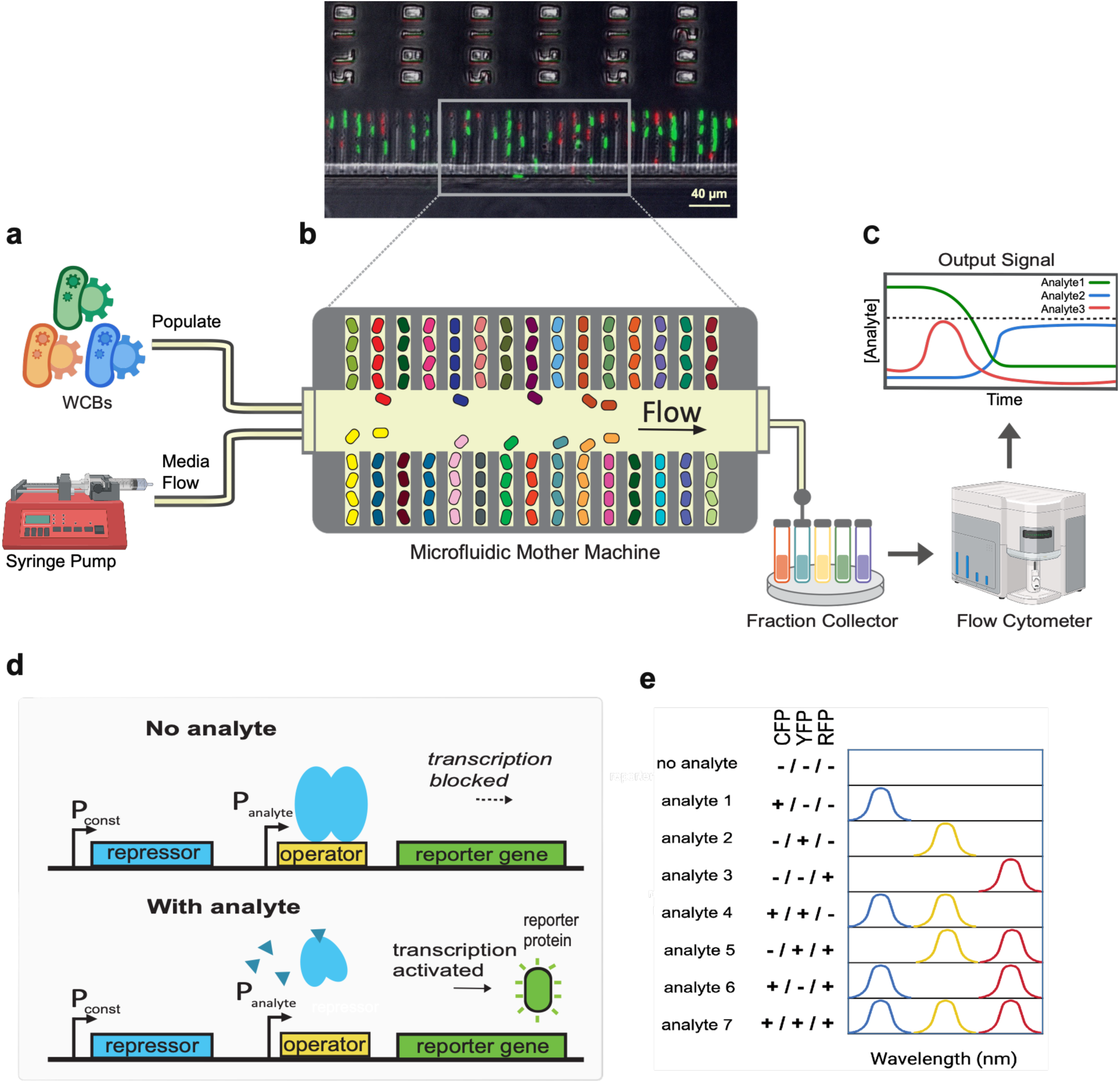
The BOSMEM platform architecture and biosensor design principles. **a,** BOSMEM deploys a genetically distinct library of engineered whole-cell biosensor (WCB) strains, populated into the microfluidic mother machine device and subjected to continuous media flow. **b,** Schematic of the microfluidic mother machine: WCB strains are loaded into individual dead-end growth channels, physically isolating each strain while continuous flow through the shared central channel sustains all strains in stable metabolic activity. Inset, representative fluorescence micrograph of biosensor strains within the device. Scale bar, 40 µm. **c,** Device effluent is collected via a fraction collector and profiled by flow cytometry, generating multiplexed analyte output signals resolved over time. **d,** Genetic architecture of the analyte-inducible reporter circuit. A constitutive promoter (P*const*) drives expression of an analyte-specific repressor that represses the transcription from an analyte-sensitive promoter (P*analyte*), silencing reporter gene expression in the absence of target analyte. Analyte binding reduces repressor-DNA affinity, activating transcription and producing fluorescent reporter protein. **e,** Combinatorial spectral encoding strategy. Each WCB strain expresses a unique combination of CFP, YFP, and RFP reporters, generating a binary activation signature across three optical channels. Seven distinguishable analyte identities are encoded using this three-fluorophore scheme, illustrating how *n* optical channels can resolve up to 2*ⁿ* − 1 distinct analytes.

## Results

### Design of Mother Machine and bacterial sensor strains

The BOSMEM platform populates a single mother machine microfluidic device with many genetically distinct WCBs simultaneously, creating a compact architecture for parallel chemical detection (Fig. 1a-c). A persistent challenge in deploying biosensor strain libraries is that open co-culture formats allow unrestricted resource competition, driving differential growth rates that shift population composition over time and erode sensing reliability^19^. The mother machine addresses this directly through its physical architecture (Fig. 1b). The device comprises a central feeding channel that delivers fresh medium continuously by pressure-driven laminar flow, and thousands of miniature dead-end growth channels branching perpendicularly from it. A cell occupying the closed end of each growth channel, termed the mother cell, divides repeatedly; daughter cells are progressively displaced toward the open end of the channel and swept away by the flow, sustaining steady-state growth and gene expression under continuous nutrient supply^20,21^. The narrow geometry (1 μm) of each growth channel physically isolates its resident strain from all others, so strains in neighboring channels share the same nutrient source but never occupy the same growth space, structurally eliminating inter-strain competition without relying on media formulation. We rationalized that this physical partitioning would be particularly important as library size grew, because strategies for managing inter-strain competition would become progressively less effective as the number of co-resident strains increased^24^. BOSMEM repurposes the mother machine architecture as a multiplexed biosensing platform by loading a distinct WCB into each channel, converting thousands of isolated microchambers into an array of independent, continuously refreshed sensor units operating in parallel.

Analyte detection proceeds through continuous exposure of the entire WCB library to the inlet sample stream. Analytes dissolved in the incoming medium diffuse from the main feeding channel into the growth channels, ensuring that every biosensor strain experiences the same chemical environment^25^ simultaneously, and that changes in analyte concentration at the inlet are rapidly propagated to all sensors. Each biosensor strain carries an inducible genetic circuit that, upon encountering its cognate analyte, activates expression of one or more spectrally distinct fluorescent proteins detectable by fluorescence microscopy or flow cytometry of the device effluent (Fig. 1b-c). The continuous-flow format provides a physical basis for reversible temporal detection. When a target analyte is present, the corresponding biosensor activates and sustains fluorescence signal in response to analyte concentration. When the analyte is withdrawn, fresh analyte-free medium displaces the inducer from the growth channels, reducing intracellular inducer concentration and allowing the repression of the fluorescent reporter genes to occur again, returning the biosensor to its baseline off state. BOSMEM can therefore track not only the presence but also the temporal dynamics of analyte fluctuations in a continuously monitored sample stream, without requiring manual intervention or reagent replenishment between measurements.

To maximize the number of detectable analytes within a fixed optical footprint, each WCB strain was engineered to express a unique combination of three spectrally distinct fluorescent proteins as its activation output: cyan fluorescent protein (CFP), yellow fluorescent protein (YFP) and red fluorescent protein (RFP). Each strain carries a plasmid encoding an analyte-specific transcriptional repressor and its cognate operator, placed upstream of one or more FP reporter genes (Fig. 1d). In the absence of the target analyte, the constitutively expressed repressor occupies the operator and blocks FP transcription, maintaining the biosensor in a fluorescently silent off state. Analyte binding (Fig. 1d) induces a conformational change in the repressor that reduces its affinity for the operator, granting RNA polymerase access to the promoter and initiating FP expression. The use of three spectrally distinct FPs as orthogonal output channels transforms each biosensor activation event into a binary three-bit signal, where each bit represents the presence or absence of CFP, YFP or RFP fluorescence. This combinatorial encoding scheme exponentially expands distinguishable biosensor identities relative to the number of optical channels required: a combinatorial scheme using *n* FPs yields 2^n^ − 1 unique non-zero codes, compared to at most *n* codes in a one-FP-per-strain approach. With three FPs this generates seven unique spectral codes, each mapped unambiguously to a specific target analyte, with the all-off state reserved for the no-analyte null condition (Fig. 1e).

### Biosensor strains exhibit reversible turn-on and turn-off kinetics in batch culture and within the mother machine

To confirm WCB functionality before device integration, we constructed a simplified WCB library – three single-fluorescent-protein biosensor strains (Fig. 2a) – and characterized their induction responses in batch monoculture. The three circuits used well-established transcriptional repressor systems: an IPTG-inducible *lac* repressor controlling YFP expression (IPTG→YFP), an arabinose-inducible *araC* repressor controlling CFP expression (ara→CFP) and an anhydrotetracycline (aTc)-inducible *tetR* repressor controlling RFP expression (aTc→RFP). Each strain was grown individually in M9-rich medium, induced at various concentrations of analyte (Fig. 2b), and fluorescence output was normalized to optical density at 600 nm (OD_600_). Activation times, defined as the time at which fluorescence response for induced sample differs substantially from non-induced sample (p < 0.05), varied across the three strains: 130 min for IPTG→YFP (10 μM), 50 min for ara→CFP (6700 μM) and 60 min for aTc→RFP (1 μM). We attribute the differences in activation time to variability in genetic engineering (i.e., repressor/promoter strength) and inducer concentrations utilized for the biosensors. Fluorescent protein maturation kinetics, which are known to vary across FP variants^26^, may also contribute, though we did not isolate this effect. These monoculture measurements established baseline detection thresholds and induction kinetics that served as reference points for interpreting device-level data.

**Fig. 2.**
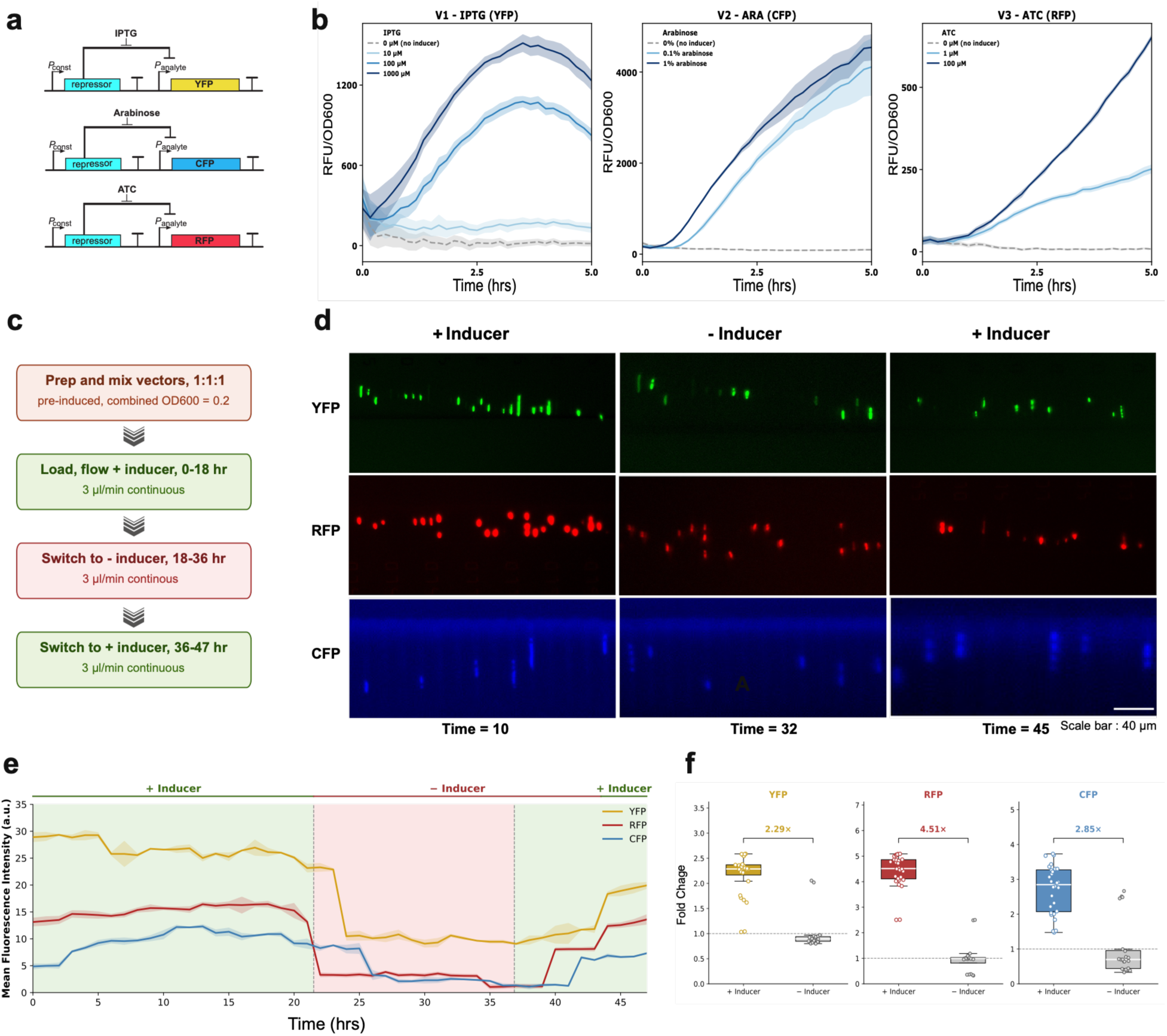
Validation of engineered biosensor functionality in batch and microfluidic culture systems. **a,** Genetic circuit schematic: a constitutively expressed repressor blocks the analyte-inducible promoter (P_analyte_) driving reporter gene expression; analyte binding relieves repression, activating transcription. **b,** Batch monoculture induction kinetics for biosensors V1 (IPTG→YFP), V2 (arabinose→CFP), and V3 (aTc→RFP) across a range of inducer concentrations, normalized to OD_600_. Shading, ± 1 S.D. (n = 3) **c,** Experimental protocol for mother machine turn-on/turn-off validation: WCBs were pre-induced (combined OD_600_ = 0.2), loaded under continuous inducer flow (0– 18 h), switched to inducer-free medium (18–36 h), then reinduced (36–47 h). **d,** Fluorescence micrographs of the three-strain co-culture within the device at representative timepoints (t = 10, 32, and 45 h) spanning the +Inducer/−Inducer/+Inducer cycle. Scale bar, 40 µm. **e,** Mean fluorescence intensity across all three channels over the 47 hours experiment, showing ON/OFF/ON switching in response to inducer withdrawal and reintroduction. Shading, ± 1 S.D. **f,** Fold change in fluorescence intensity between +Inducer and −Inducer conditions (YFP: 2.35×, RFP: 4.11×, CFP: 2.67×); points represent individual timepoint measurements.

We next sought to verify the turn-on and turn-off kinetics of the same three WCBs operating simultaneously within the mother machine device. Each strain was pre-induced in monoculture before device loading, facilitating initial visualization by fluorescence microscopy (Fig. 2d). The mother machine was populated with equal proportions of all three strains (OD_600_ = 0.2), and defined medium containing all three cognate inducers was delivered by continuous flow at a constant rate (3 μl/min) (Fig. 2c). The distinct fluorescence signals originating from each WCB was apparent during the first 18 hours of continuous flow, while all three inducers were present. Upon removal of the inducers from the flow, average fluorescence intensity in all three wavelength ranges decreased and remained at or near baseline from approximately 18 to 36 hours, confirming complete deactivation. Upon reintroduction of all three inducers at 36 hours, fluorescence intensity recovered in each wavelength range with turn-on times of approximately 4 hours (aTc→RFP), 6 hours (ara→CFP), and 8 hours (IPTG→YFP); however, by the end of the 47-hour experiment, fluorescence in each channel remained below its initial induction plateau (61.6%, 80.5%, and 73.0% of the initial maximum for CFP, RFP, and YFP, respectively), with RFP and YFP still increasing at the final measured timepoint (Fig. 2e). Quantifying the overall signal-to-background ratio across the full experiment, median fold change between +Inducer and −Inducer conditions were 2.35× for YFP, 4.11× for RFP, and 2.67× for CFP (Fig. 2f). Together, these results demonstrate that all three biosensor strains undergo reversible, analyte-dependent switching within the mother machine across at least two induction cycles over 47 hours, though full recovery to initial signal amplitude may require longer than the window monitored here.

The distinct turn-off and turn-on kinetics observed across the three biosensors likely reflect the combined influence of continuous-flow inducer displacement and the intrinsic regulatory properties of each circuit. The differences in turn-off time across strains are consistent with distinct repressor–operator affinities and inducer clearance rates specific to each system^22^. Notably, this ordering did not fully hold during reactivation, where aTc→RFP remained fastest but ara→CFP recovered before IPTG→YFP, suggesting turn-off and turn-on kinetics are not governed by a single shared mechanism, and that additional factors, such as inducer diffusion into growth channels or reporter maturation kinetics, may differentially affect activation versus deactivation. Kinetics in the mother machine may differ from those seen in batch culture due to slower growth rates. Interestingly, removal of the analyte from the mother machine displays delayed reaction followed by a sharp decrease in YFP and CFP signal, as opposed to a steady reduction in signal. These results could indicate the bottleneck for biosensor deactivation, where reduction in signal relies on dilution of fluorescent proteins through cell division rather than fluorescent protein degradation. Slower growth rates in the mother machine could lead to the observed delays in signal reduction. These results establish that the mother machine supports dynamic, largely reversible analyte sensing on a timescale of hours, governed by the physical properties of the continuous-flow format together with the intrinsic regulatory characteristics of each biosensor circuit.

### A seven-member biosensor library responds to chemically diverse analytes through distinct three-channel spectral codes

Having confirmed individual biosensor functionality and reversible switching kinetics, we sought to construct a full seven-member WCB library that exploits the combinatorial spectral coding scheme across three fluorescent proteins. Seven *E. coli* strains (B1 through B7) were engineered to respond to chemically diverse small molecules spanning inorganic heavy metals, aromatic acids, aldehydes, a monosaccharide and a tetracycline derivative, collectively demonstrating the breadth of analyte classes compatible with the combinatorial^27^ spectral coding scheme (Fig. 3a). Each strain was designed to produce a unique combination of YFP, CFP, and RFP outputs upon induction by its cognate analyte, generating a three-bit spectral code that distinguishes it from all other library members. The seven target analytes were cuminic acid (B1, cym→YFP), arabinose (B2, ara→CFP), vanillic acid (B3, van→RFP), Hg^2+^ (B4, mer→YFP+RFP), formaldehyde (B5, form→YFP+CFP), anhydrotetracycline (B6, aTc→CFP+RFP) and acrylic acid (B7, acr→YFP+CFP+RFP). Each biosensor was tested with a range of analyte concentrations (Fig. 3c-d). Results were used to determine detection limits (Fig. 3d) and activation times (Fig. 3c) for each biosensor. The wide range of detection limits for the biosensors (0.0023 μM for B5 - 100 μM for B7) is likely a symptom of variability in affinity constant between repressors and their cognate inducer molecules, though other genetic design factors may play a role. Interestingly, biosensors that produced RFP output (B3, B4, B6, B7) had longer activation times, which may be attributed to higher maturation half-time of RFP as compared to YFP and CFP.

**Fig. 3.**
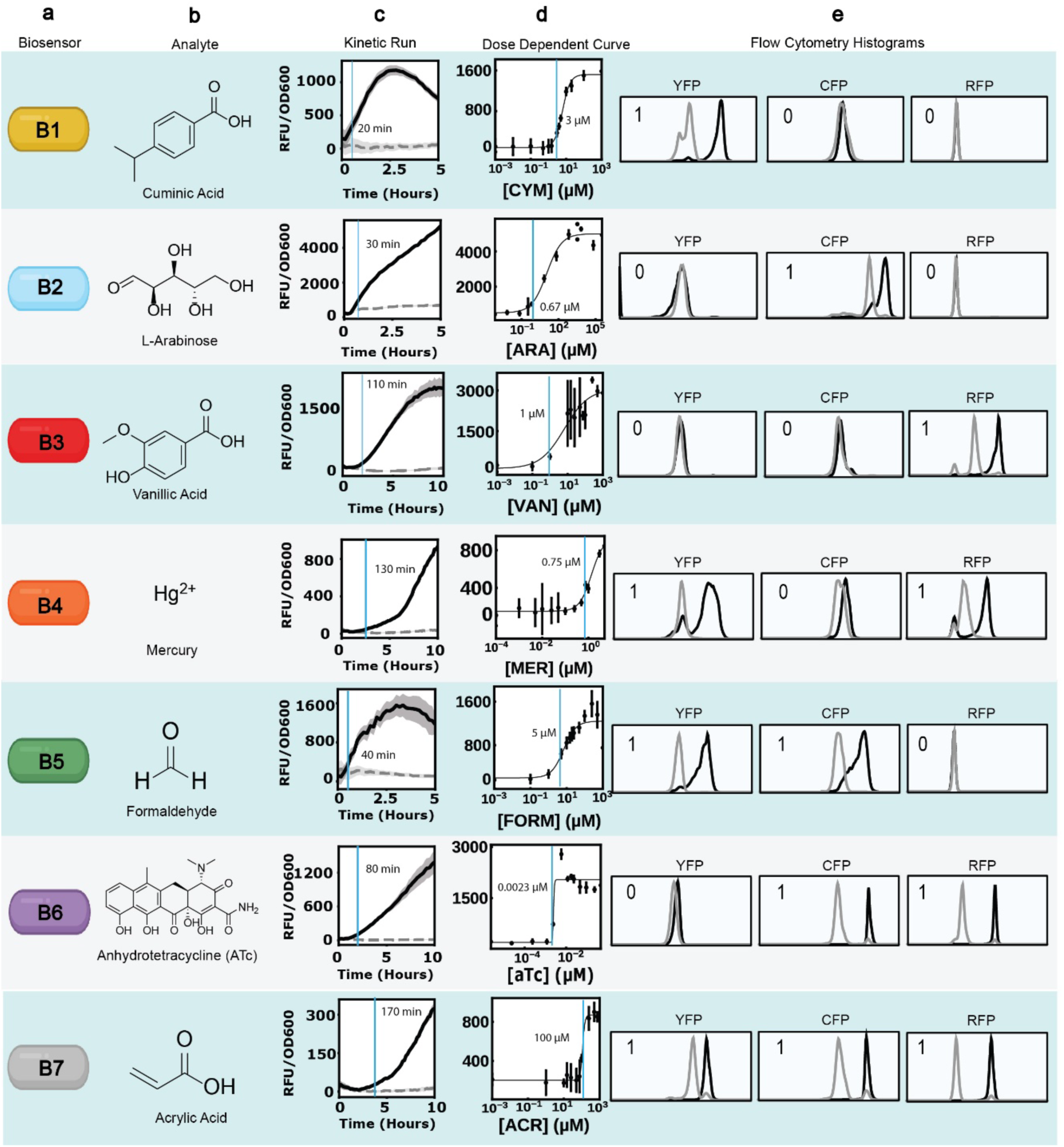
Engineering and validation of seven multiplexed biosensors. Columns display different characteristics for the biosensors. **a,** Biosensor identification code. **b,** Analyte structure. **c,** Normalized kinetic runs for each biosensor (B1 – YFP Signal; B2 – CFP Signal; B3 – RFP Signal; B4 – RFP Signal; B5 – YFP; B6 – RFP; B7 - RFP) with activation time noted by the blue line. Non-induced samples are represented by dotted gray line, while induced samples represented by black solid line. Shading, ± 1 S.D. (n = 3) **d,** Dose dependent curves for each biosensor (B1 – YFP Signal; B2 – CFP Signal; B3 – RFP Signal; B4 – YFP Signal; B5 – YFP; B6 – CFP; B7 - YFP) with detection limit (LOD) noted by blue line. (n = 3) **e,** Flow cytometry histograms for each biosensor monoculture. Non-induced samples are represented by gray line, while induced samples are represented by black lines. Detection limits and activation times for dual-channel sensors were determined by FP output that produced highest LOD concentration or activation time.

The spectral code of each strain was validated by flow cytometry after 18 hours of induction in conventional tube culture. Non-induced monoculture populations served as baseline controls, and induced monoculture populations were analyzed in parallel to quantify the shift in fluorescence intensity in each channel (Fig. 3e). To quantify the degree of phenotypic heterogeneity in each activated biosensor population, we assigned each a robust coefficient of variation (rCV) (Table S3). Single-FP sensors produced clean, well-resolved population shifts exclusively in their designated output range. B1 (cym→YFP) produced a rightward shift in the YFP range with an >87-fold median increase in YFP signal compared to its non-induced sample. B2 (ara→CFP) and B3 (van→RFP) showed equivalent behavior in their corresponding wavelength ranges, with a >64-fold (CFP) and >44-fold (RFP) median signal increase, respectively. For all three biosensors, the induced and non-induced populations were well separated, confirming low basal expression and high signal-to-noise ratio across all single-channel biosensors.

Dual-FP sensors demonstrated simultaneous population shifts in both designated wavelength ranges with no appreciable signal in the unassigned range. B4 (mer→YFP+RFP), induced with 3 μM mercury, produced coincident YFP (>34-fold median increase) and RFP shifts (>27-fold median increase) while the CFP channel remained at baseline. The YFP shift in B4 was notably broader (rCV = 152) than in the single-channel YFP sensor B1, suggesting population heterogeneity in the mercury-sensing response at the tested concentration, while the activated RFP population showed more homogeneity (rCV = 93.6). B5 (form→YFP+CFP), induced with 25 μM formaldehyde, showed a clean YFP shift (>30-fold median increase) accompanied by a distinct rightward CFP shift (>15-fold median increase) with no signal observed in the RFP range. B6 (aTc→CFP+RFP), induced with 0.023 μM anhydrotetracycline, produced a well-separated RFP shift (>192-fold median increase) and a smaller but distinct CFP shift (>51-fold median increase), consistent with differential expression levels driven by the aTc-responsive circuit at the tested concentration. The triple-FP sensor B7 (acr→YFP+CFP+RFP), induced with 100 μM acrylic acid, produced simultaneous shifts in all three wavelength ranges, completing the seven-code combinatorial space (YFP >6-fold; CFP >31-fold; RFP >262-fold median increase) (Fig. 3e).

Across all seven strains the induced populations were distinguishable from their non-induced counterparts in the relevant wavelength ranges. While most strains did not show spurious signal in off-target channel (Fig. 3e), one within-channel exception is worth noting: B2’s (ara→CFP) non-induced population showed a low-level shoulder in the CFP channel, consistent with basal leakiness of the araC promoter system reported previously^28^. This leaky subpopulation was well below the induced population in CFP intensity and was separable by gating, as demonstrated in the multiplexing analysis below (Fig. S10). Each of the seven strains therefore occupies a spectrally distinct position in the three-dimensional YFP/CFP/RFP fluorescence space, providing the unique spectral codes required for unambiguous analyte identification within a mixed co-culture.

To assess whether the seven spectral codes remained distinguishable when all biosensor strains were present simultaneously, we developed a flow cytometry gating strategy for analyte identification within a complex co-culture. Induced monocultures of each strain were prepared separately, washed and combined at equal densities to generate a reference co-culture sample in which all seven biosensors were activated (Fig. S3). The multiplex-ability of the platform relies on the unique spectral output produced by each activated biosensor. Sequential two-dimensional gating through the YFP, CFP, and RFP scatter plots was used to isolate individual strain populations.

We utilize B6 (aTc→CFP+RFP) gating isolation as the primary demonstration example because its CFP⁺ YFP⁻ RFP⁺ code requires two sequential gating steps and therefore illustrates the full resolution capacity of the strategy. In the co-culture sample with all biosensors activated, an initial gate selecting CFP⁺ YFP⁻ cells identified a population containing both B2 (ara→CFP, which is CFP⁺ RFP⁻) and B6 (aTc→CFP+RFP, which is CFP⁺ RFP⁺) (Fig. S3). The selected population was carried forward to a second scatter plot in which two distinct sub-populations were visible. Selecting the RFP⁺ sub-population within this gate isolated B6, consistent with its expected spectral code. The same sequential gating procedure was applied to a parallel co-culture sample in which only B6 was activated, yielding an isolated population whose scatter profile closely matched the B6 population extracted from the fully activated co-culture. An RFP⁻ sub-population was also visible in the B6-only sample at the first gating step, indicated by the gray arrow (Fig. S3). This population likely represents leaky CFP expression from uninduced B2, consistent with the basal araC promoter activity observed in monoculture (Fig. 3e, Fig. S10). Overlay of the B6 populations isolated from both gating conditions confirmed high spatial correspondence across all three FP channels between co-culture with all biosensors induced and co-culture with only B6 induced. Thus, B6 can be accurately identified regardless of whether the surrounding co-culture contains one or all seven activated strains.

These results demonstrate that the three-channel combinatorial spectral coding scheme supports reliable identification of individual biosensor populations within a seven-strain co-culture by sequential flow cytometry gating, without requiring physical separation of strains. The gating strategy was successfully extended to all seven library members (Fig. S4–S9), establishing a generalizable framework for analyte identification that scales with the combinatorial code space rather than with the number of physically distinct biosensor reservoirs.

### The BOSMEM platform sustains stable biosensor population ratios and analyte-responsive spectral outputs over several days

A fundamental requirement for a continuous chemical monitoring platform is the ability to maintain both stable population composition and reliable sensing function over extended periods. To evaluate these properties, we conducted a multi-day longevity experiment in which all seven biosensor strains were maintained either in batch co-cultures or simultaneously within the mother machine, with distinct analytes introduced sequentially on designated days. Analyte-response fluorescence output was tracked daily by flow cytometry over an eight-day induction schedule, with relative cell densities quantified across the YFP, CFP, and RFP channels and their double- and triple-positive combinations. Population composition was assessed separately by qPCR over a nine-day schedule using the same sequential induction design. This allowed us to assess whether each biosensor strain retained its expected spectral response after multiple days of co-culture, and whether population proportions remained sufficiently stable.

In batch culture, flow cytometry histograms across the YFP, CFP and RFP channels showed progressive deterioration of WCB functionality over the eight-day experiment (Fig. 4b). Batch culture experiments were conducted by combining the seven biosensors into a co-culture at equal ratios (Fig. 4a). Each day, a co-culture aliquot was collected for flow cytometry analysis, and cells were “passaged” to renew media as well as wash away analytes introduced on previous days. On each day, flow cytometry histograms displayed a single narrow peak corresponding to the dominant uninduced population. Induction of B1 with cuminic acid on Day 1 resulted in a YFP-only shift aligning with expected signal that was generated from B1 activated monoculture. On Day 4, following induction of B2 with arabinose, the expected CFP-only signal was generated by a smaller cell population as compared to results on day 2. By Day 8, following induction of B3 with vanillic acid, we found that the expected population of RFP-only cells was negligible. To determine whether WCB deterioration in batch cultures is driven by population distribution instability, we conducted qPCR experiments on batch cultures containing all seven biosensors over a separate nine-day experiment. We designed primer set specific for each biosensing plasmid and converted Ct values from qPCR experiments into relative cell density (RCD) for each biosensor. From this, we calculated a differential cell density value (DCD) to visualize population drift of the biosensors. This analysis confirmed progressive compositional drift over the nine-day tube culture period, with B4 increasingly dominating the culture at the expense of all other biosensors (Fig. 4e).

**Fig. 4.**
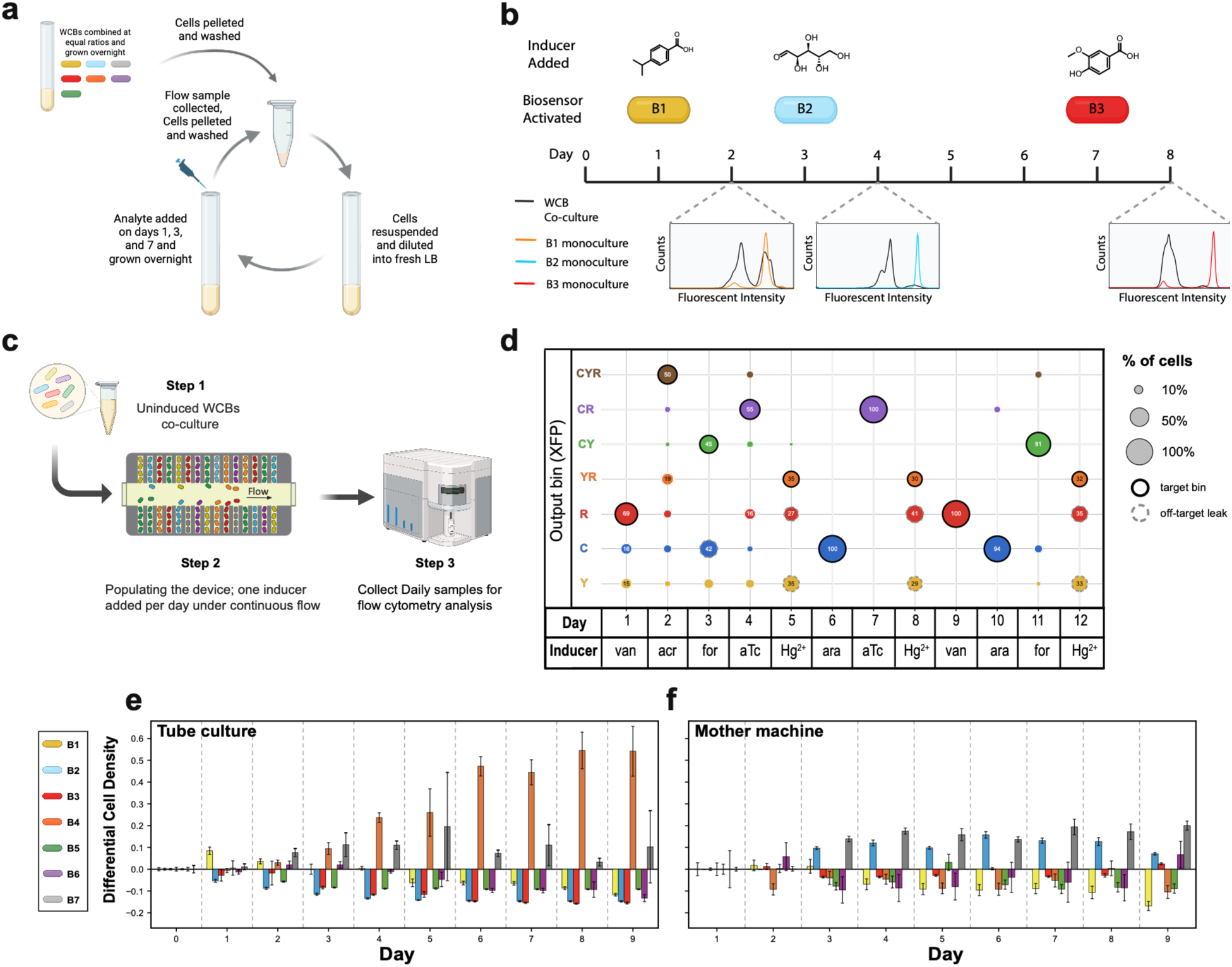
Longevity and population stability of WCBs in the BOSMEM platform. **a**, Workflow for the batch-culture longevity experiment: seven WCB strains were co-cultured at equal starting ratios, sequentially induced with strain-specific analytes on designated days, and daily flow cytometry samples were collected after washing and resuspension in fresh medium. **b**, Sequential single-analyte induction schedule and validation in batch co-culture. Analytes activating B1, B2, and B3 (chemical structures shown) were introduced on Days 1, 3, and 7, respectively; inset flow cytometry histograms compare the seven-strain co-culture (black) to the corresponding activated monoculture (colored) at each induction timepoint, confirming that co-culture fluorescence shifts match the expected monoculture signal. **c**, Workflow for the mother machine longevity experiment: uninduced WCBs were co-cultured, loaded into the device under continuous flow with one inducer introduced per day, and daily effluent samples were collected for flow cytometry analysis. **d**, Output bin composition across the 12-day sequential induction schedule within the mother machine, with bubble size representing the percentage of cells occupying each spectral bin (C, Y, R, and their double- and triple-positive combinations) on each day; bold outlines denote the expected target bin for that day’s inducer, dashed outlines denote off-target leakage. (**e, f**) Differential cell density (relative to Day 1) for each biosensor strain (color key, left) over the course of the longevity experiment in tube culture (n = 3) (**e**) versus the mother machine (n = 3, except B6 Day 1 RCD n = 2) (**f**), showing progressive compositional drift in batch co-culture contrasted with stable population proportions in the device.

The mother machine produced a qualitatively distinct result. Fig 4d plots the measured percentage of cells occupying each non-null spectral bin across every day of the experiment, with bold outlines marking the bin expected to respond to that day’s inducer. Each unique FP combination represents a bin (e.g., CFP^-^/YFP^+^/RFP^+^ bin, or CFP^-^/YFP^+^/RFP^-^ bin). On Day 1, induction with vanillic acid produced a CFP^-^/YFP^-^/RFP^+^ signal (i.e., 69% of cells collected in the MM effluent fraction for Day 1 were in the RFP^+^ bin) consistent with the expected B3 (van→RFP) response, while the CFP^-^/YFP^+^/RFP^-^ and CFP^+^/YFP^+^/RFP^-^ bins remained close to 0% populated. On Day 2, induction with acrylic acid produced a population in the expected triple-positive YFP^+^CFP^+^RFP^+^ bin, with the YFP^+^CFP^+^RFP⁺ combination rising to a level consistent with B7 activation. Across all twelve days of the experiment, the bin with the largest measured population matched the expected target bin for that day’s inducer (bold outline, Fig. 4d), confirming that the correct biosensor responded to each inducer at each timepoint. However, we did observe a few off-target events: on Day 3, following formaldehyde induction, and Days 5, 8, and 12 following mercury induction. We think these unexpected results most likely arise from genetic instability of plasmid-based WCB engineering, and we address this in more detail in the Discussion section. We used the same qPCR-based protocol to determine time-dependent variation in relative population densities in the mother machine as we did for batch co-culture experiments. Overall, we observed substantially lower variability in the differential cell densities of the seven biosensors over time in the mother machine as compared to batch co-culture (Fig. 4e, f). This likely arises from the mother machine channel structure, which is designed to minimize resource competition between mother cells, allowing for parallelized single-cell tracking and experimentation. These results demonstrate that the mother machine architecture maintains both population stability and sensing accuracy across a multi-day sequential detection schedule that would be incompatible in a co-culture format without similar single cell confinement.

The contrast between tube culture and mother machine performance under identical induction schedules directly validates the central design principle of BOSMEM. In batch co-culture, unrestricted inter-strain resource competition produces population composition drift on a timescale of days, which in turn corrupts the spectral code assignment by changing the relative fluorescence contributions of each strain to the mixture signal^29^. The mother machine eliminates this drift mechanism by physically partitioning each strain into its own set of growth channels under continuous nutrient flow, maintaining every strain at steady-state growth rate and stable ratios throughout the monitoring period^19,20^. The sustained analyte-responsive spectral outputs, combined with the stable population proportions between induction events, confirm that BOSMEM is capable of continuous, reliable multiplexed chemical sensing over 12 days.

### The BOSMEM platform generates accurate responses from analytes in a complex sample matrix

Once we confirmed BOSMEM longevity and population stability of the biosensor library, we sought to display reproducible functionality for detection in complex matrices. The common baking ingredient vanilla extract consists of an assortment of volatile compounds, alcohols, and the flavor agent vanillic acid. To display reproducible detection of vanillic acid in vanilla extract, we first confirmed B3 activation upon vanilla extract exposure with overnight induction of batch monocultures (Fig. 5a-c). Interestingly, introduction of vanilla extract at volumes 20 and 40 μl produced the highest normalized fluorescence intensity, whereas 110 μl produced a lower intensity which could be a result of cell malfunction from increased concentrations of incompatible extract components.

**Fig. 5.**
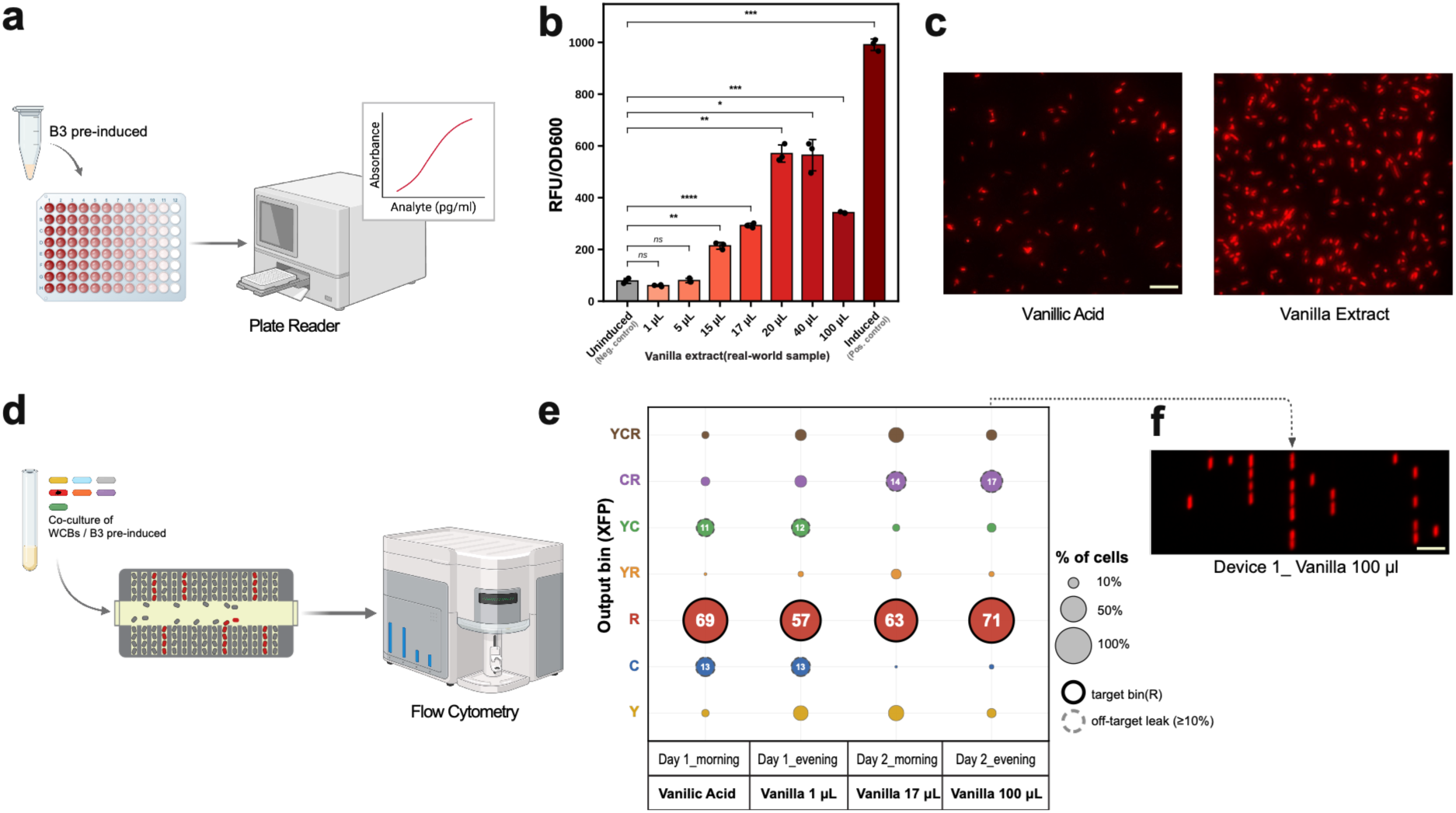
Application of BOSMEM to sense analytes withing a complex matrix: detection of vanilla extract by biosensor B3. **a**, Workflow schematic for dose-response quantification: B3 culture was aliquoted into a 96-well plate for plate-reader fluorescence measurement, generating a dose-response curve relating fluorescence to analyte concentration. **b**, Dose-dependent fluorescence response of B3 to increasing volumes of vanilla extract, normalized to OD₆₀₀. “Uninduced” denotes the no-analyte negative control; “Induced” denotes the pure-vanillic acid positive control. Asterisks denote statistical significance relative to the uninduced control (Welch’s *t*-test vs. Uninduced, Holm–Bonferroni corrected; *P < 0.05, **P < 0.01, ***P < 0.001, ****P < 0.0001; ns, not significant); error bars, ± 1 S.D. (n = 3 biological replicates). **c**, Representative fluorescence micrographs of B3 induced with pure vanillic acid (left) versus vanilla extract (right), confirming that the biosensor responds to the analyte within a complex, unpurified matrix. Scale bar, 20 µm. **d**, Workflow for mother machine-based detection of vanillic acid within vanilla extract: a co-culture of all seven WCB strains was loaded into the device and profiled by flow cytometry at morning and evening timepoints over two days, with vanillic acid or vanilla extract introduced before each sampling point. **e**, Output bin composition (percentage of cells in each spectral bin, C, Y, R and their double- and triple-positive combinations) across four sequential timepoints – Day 1 morning, Day1 evening, Day 2 morning and Day 2 evening – following introduction of vanillic acid and increasing volumes of vanilla extract (1, 17, and 100 μ, respectively) into the mother machine. Bold outlines denote the expected target bin (R); dashed outlines denote off-target leakage (≥10% of cells). **f**, Representative fluorescence micrograph corresponding to the Day 2 evening (100 μ vanilla extract) timepoint in (e), showing RFP⁺ cells within mother machine growth channels. Scale bar, 20 µm.

To confirm reproducible detection of vanillic acid from vanilla in the BOSMEM platform, we conducted an experiment housing the seven established biosensors in the mother machine, followed by introduction of increasing volumes of vanilla extract (Fig. 5d). Similar to the BOSMEM longevity experiment, time points were analyzed using multi-step gating strategy in flow cytometry. Upon exposure of the BOSMEM platform to vanillic acid on Day 1 (morning), the biosensors maintained their expected spectral response with a 69% population within the R (RFP+) bin. A consistent response was displayed upon introduction of 1 μl of vanilla extract (Day1, evening) into the system (57% RFP+ population) and maintained with treatment of increasing volumes of vanilla (63% for 17 μl; 71% for 100 μl) (Fig. 5e-f). We also noted off-target signal exceeding our 10% leakage threshold in two bins across the experiment: C and YC at Day 1 (13% and 11-12%, respectively) and CR at the two highest vanilla volumes (14% at 17 μl, 17% at 100 μl).

As a whole, these results demonstrate that BOSMEM correctly identifies its target analyte against the dominant background bin within a complex, real-world matrix, while also revealing modest off-target activity that will need to be characterized before the platform is deployed on unpurified samples in the field.

## Discussion

Here, we developed the BOSMEM detection platform by housing multiplexed WCBs in the mother machine microfluidic device. BOSMEM displayed turn-on/off detection capabilities with a library of biosensors over the course of several hours. We used combinatorial fluorescent protein outputs to create a library of seven WCBs that could all be uniquely distinguished from one another with single-cell detection techniques. We employed a multistep gating strategy to identify activated populations in longevity tests, demonstrating that the biosensors can maintain functionality over the course of 12 days, longer than typical reported mother machine experiments, which are commonly run over hours to a few days^20,21,30,31^. Additionally, population distribution experiments showed that the BOSMEM platform leads to higher co-culture population stability as compared to batch culture conditions, suggesting that mother machine-like devices facilitating the long-term culture of engineered microbes could be valuable platforms biotechnologies.

Previous studies have demonstrated many materials- and device-based approaches to WCB sensing platforms^32,33,34^ that are more amenable to field deployment, but they largely focus on one-time use and small numbers of analytes. In contrast, BOSMEM is designed to extend toward time-resolved outputs with tens-to-hundreds of unique biosensors. This approach is enabled by the continuous media flow and cell growth supported by the compact mother machine design. The specific analytes that we targeted were based on the ready availability of well-characterized inducible repressor genes. However, the identity of analytes that the system responds to could be altered easily with the very large number of known inducible transcription factors^12,35,36^. Because we chose to combine three fluorescent proteins to create our multiplexed output signal, we were limited to a seven-member library of WCBs. It may be possible to use additional outputs, including additional fluorescent proteins or hyperspectral signals^37^ to accommodate larger WCB libraries, though concerns about spectral overlap and the metabolic burden of producing so many FPs simultaneously may have to be addressed. Future iterations of BOSMEM may benefit from other approaches to single-cell detection, like automated fluorescence microscopy, or electrochemical outputs.

The time-resolved output of BOSMEM was limited in resolution by the frequency of fraction collection from the effluent of the mother machine. We found that higher frequencies led to fractions with insufficient cell counts to complete flow cytometry analysis. Thus, the time resolution of BOSMEM could be enhanced significantly by in situ monitoring of WCB activation inside the mother machine. Notably, several previous mother machine papers successfully monitored hundreds of unique mother cells with custom automated microscopy and image analysis protocols^38,39,40^. The OFF-to-ON kinetics varied widely between WCBs but were limited to at least 2 hours on the faster end. This likely approaches a lower limit since transcription, translation, and protein folding must take place between analyte introduction and signal generation. This transition speed might be improved by genetic design – for example, WCBs that use riboswitches or allosteric proteins as transduction elements, skipping over some steps of the central dogma^41–43^. The ON-to-OFF transition speed of the WCBs may have been limited by the rate of fluorescent protein degradation or the rate of cell division. Future studies may investigate enhancing cell growth rate or reporter protein degradation to shorten this transition time.

We performed extensive quality control experiments to ensure that each WCB in our library performed as designed in tube culture immediately prior to populating the mother machine (Fig. S2). Despite this, we found that long-term co-culture experiments in the mother machine led some of the WCBs to exhibit unexpected behaviors. On Day 3, the formaldehyde-responsive WCB (B5) showed erroneous population of the “C” bin in addition to the expected “YC” bin. However, on Day 11, it showed a more expected response (81% in the “CY” bin). In contrast, the mercury ion sensor (B4) gave the same erroneous response each of the three times it was induced (Day 5, 8, and 12). There may be many possible explanations for these aberrant results. We think it is unlikely that the signal in the aberrant bins arises from the activation of an off-target WCB. Identifying genetic mutations to the plasmids encoding the biosensor elements may be able to explain these results, though a rigorous investigation may require a combination of continuous fluorescence microscopy, cell sorting, and genetic sequencing of many cell sub-populations. Nevertheless, the fact that B4 malfunctioned in the same manner each time it was induced suggests that aberrant WCB behavior may still be useful in the context of identifying individual input analytes as long as it remains consistent.

Overall, we envision the advancement of BOSMEM along several axes. For example, WCB performance could be improved through more sophisticated genetic engineering or better cell-device interfaces. The deployability of BOSMEM may be enhanced through custom device engineering and modern advances in optical detection techniques that might facilitate in situ single-cell monitoring. As such, we view BOSMEM as a promising platform for further development toward olfactory-like continuous environmental monitoring.

## Methods

### General reagent, instrument, and resources

M9 minimal salts, calcium chloride, and magnesium sulfate were purchased from Sigma-Aldrich. LB broth and casamino acids were purchased from Research Products International (RPI). Agar was purchased from Fisher Scientific. Trace elements and carbenicillin were purchased from Teknova. Glucose was purchased from VWR Chemicals. Black, clear-bottom well 96-well plates were purchased from Greiner. qPCR well plates were purchased from VWR.

All primers utilized in this work were purchased from Genewiz. Q5 PCR master mix, Gibson assembly master mix, site-directed mutagenesis kit, Luna qPCR master mix, and BL21 (DE3) chemically competent cells were purchased from New England Biolabs (NEB). Mach1 cells were purchased from Thermo Fisher. Gene blocks were ordered from Twist Bioscience. Plasmid purification was done with Qiagen Miniprep kit.

Equipment utilized in this work includes: fluorescent microscope (Echo), plate reader (Molecular Devices - Spectramax M5), flow cytometer (Beckman Coulter Analyzer), thermocycler (ProFlex PCR System, Thermo Fisher), RT-thermocycler (C1000 Touch Thermal Cycler, Bio-Rad), and centrifuge (Eppendorf).

### Cell culture and induction conditions

Transformed BL21(DE3) biosensor cells were grown and induced in these conditions unless stated otherwise. Starter cultures (5 ml) were grown overnight in a shaking incubator at 37°C and 225 rpm with carbenicillin (100 μg/ml). Starter cultures were diluted the next day (1:100) into fresh media, and grown until cells reached OD of 0.4-0.6. Cultures were then induced with corresponding inducers listed below:

B1 – 10 μM cuminic acid

B2 – 6700 μM L-arabinose

B3 – 100 μM vanillic acid

B4 – 3 μM mercury chloride

B5 – 25 μM formaldehyde

B6 – 0.023 μM anhydrotetracycline

B7 – 100 μM acrylic acid

### Construction of strains and plasmids

Plasmids were constructed using molecular cloning techniques. Vector and gene fragments were amplified via PCR or purchased through custom gene block orders. Plasmid formation was conducted via Gibson assembly or site directed mutagenesis for point mutations. Following assembly, plasmids were transformed into Mach1 cells and plated on LB agar containing carbenicillin (100 μg/ml). Colonies were selected and cultured for plasmid purification followed by whole-plasmid sequencing through Plasmidsaurus. Plasmids verified to have expected sequencing were transformed into BL21 (DE3) cells (New England Biolabs) for biosensor testing.

Plasmids containing ptrc99a backbone, CFP, YFP, and mCherry gene were gifted from Elizabeth Libby. Plasmids were purchased from Chris Voigt depository on Addgene for certain repressor, transporter, and promoter sequences (*cymR*-AM – #108525; *vanR*-AM - #108527; *araE* - #108530; *acuR* - #108536). Plasmid encoding *frmR* repressor and promoter sequence was purchased from David Liu depository on Addgene (#104068). DNA fragments encoding repressor/promoter pairs *araC, tetR, lacI,* and *merR* were amplified from plasmids prepared in our previous work. Antibiotic resistance genes were also amplified from plasmids prepared in our previous work.

### Fabrication of the microfluidic devices

The microfluidic mother machine design was based on a mold originally developed by Wang et al.^20^, obtained as a PDMS replica (gift from S. Jun) that was cast in epoxy to generate a durable master. The device contains hundreds of dead-end lateral growth channels (∼1 μm × 1 μm cross-section, 25 μm length) branching off a central flow channel (25 μm high, 100 μm wide). PDMS chips were cast from the epoxy master following the general approach described previously^44^: a 10:1 (base: curing agent) PDMS mixture was poured over the mold and cured at 65°C for 90 min. After curing, chips were peeled from the master, and inlet/outlet ports were punched using a 0.75 mm biopsy punch (Harris Uni-Core, WPI). Chips were bonded to glass coverslips by oxygen plasma treatment (30 W, 10 s; Diener plasma etcher), as described previously^44^. Because plasma treatment transiently renders both surfaces hydrophilic, devices were backfilled with 2 μL of 50 mg/mL bovine serum albumin within 5 min of bonding and incubated at 37°C for 30 min to passivate internal channel surfaces and reduce nonspecific cell adhesion.

### Fluorescence kinetic and dose dependent data for all biosensors

Cells were grown, diluted into M9 media, induced, and plated in 96-well plate for overnight plate reader fluorescence reading. Plate reader was set to read CFP (ex: 433 nm, em: 475 nm), YFP (ex: 513 nm, em: 530 nm), and RFP (ex: 587 nm, em: 610 nm) relative fluorescence units (RFU) as well as OD600 every ten minutes overnight. RFU values were normalized by OD600 per sample.

Dose dependent data were collected at maximum RFU/OD600 values within the initial plateau for each biosensor tested at different concentrations. These values were plotted against concentrations tested to generate final dose dependent curves. Detection limit was determined based on the lowest concentration tested that generated a significant RFU/OD600 value as compared to non-induced sample.

### Turn-on/turn-off kinetics in the mother machine

The three single-fluorescent-protein biosensor strains were named consistent with the B1–B7 nomenclature used elsewhere in this study: V1 (IPTG→YFP), V2 (ara→CFP), and V3 (aTc→RFP). Each strain was grown to exponential phase in M9 defined medium and pre-induced in monoculture for 6 hours with their cognate inducer (10 µM IPTG, 0.1% w/v L-arabinose, or 40 ng/mL anhydrotetracycline, respectively) prior to device loading, to enable initial visualization by fluorescence microscopy. Equal proportions of the three strains were combined to a final OD₆₀₀ of 0.2 and loaded into the mother machine device as described above. Following loading, M9 defined medium containing all three inducers (10 µM IPTG, 0.1% w/v L-arabinose, 40 ng/mL aTc) was delivered by continuous flow at a constant rate of 3 μl/min using a syringe pump (New Era Pump Systems, Inc.). Cells were maintained at 37°C throughout the experiment. Time-lapse fluorescence microscopy was performed on an Echo Revolve microscope (model RVL-100-M; Discover Echo Inc.) equipped with a 20× objective and a Point Grey CM3-U3-50S5M camera. Images were acquired hourly across CFP, YFP, and RFP channels throughout the 47-hour experiment; at each timepoint, three images spanning the device were captured, and mean fluorescence intensity was calculated across the three images for each biosensor channel using ImageJ. Background subtraction and downstream quantification were performed in MATLAB. After 18 hours of continuous induction, the inducer-containing medium was replaced with inducer-free M9 medium of identical composition to initiate deactivation; all three inducers were reintroduced at the original concentrations at 36 hours to assess reactivation. For each channel, fluorescence values were normalized to the maximum intensity observed during the initial induction period (0–18 h) to calculate percent recovery upon reinduction. Turn-off and turn-on times were defined as the time required for average channel fluorescence to fall to or rise to [threshold, e.g., 50%] of the initial induction plateau.

### Validation of seven biosensor functionality in flow cytometer

Cells for each biosensor were grown and induced overnight. Following overnight induction, cells were washed, resuspended, and diluted to OD ∼0.2 in 1x PBS. Cells were analyzed via flow cytometry (Beckman Coulter CytoFLEX). Data were analyzed on FlowJo.

### Multiplexed tube culture analysis

Cells for each biosensor were grown and induced overnight. Following overnight induction, cells were washed, resuspended, and diluted to OD 0.2 in 1x PBS. The biosensors were combined in a culture tube at equal ratios followed by analysis on the flow cytometer. Data were analyzed on FlowJo.

### Flowjo analysis

Compensation was performed on flow cytometry data using cells producing highest YFP, CFP, RFP signals from experiment as well as BL21 cells containing SHAM plasmid as negative control. Samples were gated to exclude debris and doublet populations. Robust CV values for non-induced and induced monocultures of multiplexed biosensors were generated. Median values for non-induced and induced monocultures of multiplexed biosensors were generated to calculate fold median change.

### Mother Machine data analysis for longevity experiment

Flow cytometric events were computationally processed using a standardized gating and fluorescence-classification workflow. Events were first gated on forward and side scatter (FSC-A, SSC-A), retaining the 5th–95th percentile range to exclude debris and non-cellular events. A subsequent singlet gate was applied to the FSC-H/FSC-A ratio, retaining the 10th–90th percentile range to exclude doublets and aggregates. Spectral compensation was performed using single-color controls for the CFP, YFP and RFP channels. Spillover coefficients were estimated from the brightest 30% of events in each single-color control, after subtracting mean background fluorescence measured in the gated unstained control. The resulting spillover matrix was inverted to generate a compensation matrix, which was applied to the gated unstained control and from all 12 days prior to threshold setting and classification. Channel-specific positivity thresholds were defined as the 99th percentile of each channel’s distribution in the compensated unstained control. Fluorescence classification used a Boolean thresholding framework across the three detection channels, partitioning each gated, singlet-filtered event into one of eight mutually exclusive states: three single-positive (Y, C, R), three double-positive (YC, YR, CR), and one triple-positive (YCR). For each sample, the abundance of each state was calculated as a percentage of that sample’s total gated, singlet-filtered population; no subsampling or event-count equalization was performed across samples. The dominant fluorescence phenotype per sample was defined as the non-negative state with the highest percentage; states within 3.0 percentage points of the maximum were called co-dominant. Mean fluorescence intensity per channel was calculated from the compensated, background-subtracted values.

### Biosensor longevity test in batch culture

Starter cultures for each biosensor were grown overnight, diluted in LB (1:100), and grown until OD ∼0.4. Cells were then combined at equal ratios into a culture tube to make seven biosensor co-culture. Co-culture was grown over the course of the eight-day experiment in shaking incubator (37°C, 225 rpm). Each morning, co-culture was “passaged” through washing and resuspending in fresh LB to remove old media and analytes that were previously in the sample. The co-culture was induced on evenings on Day 1 (cuminic acid), Day 3 (L-arabinose), and Day 7 (vanillic acid). Samples were collected, washed with 1x PBS, pelleted, and stored at 4°C for later testing. At the end of the experiment, pelleted samples were resuspended in PBS and analyzed via flow cytometry. Data were analyzed on FlowJo.

### Biosensor longevity test in mother machine

Starter cultures for each of the seven biosensors were grown overnight, diluted in LB (1:100), and grown until OD₆₀₀ ∼0.4. Cells were combined at equal ratios to generate a seven-biosensor co-culture and loaded into the microfluidic mother machine device at a combined OD₆₀₀ of 0.2. Defined medium was delivered by continuous flow at 3 µl/min throughout the twelve-day experiment. A single inducer was introduced into the flowing medium each day, following the sequence: vanillic acid (100 µM, Day 1), acrylic acid (100 µM, Day 2), formaldehyde (25 µM, Day 3), anhydrotetracycline (0.023 µM, Day 4), mercury chloride (3 µM, Day 5), L-arabinose (6700 µM, Day 6), anhydrotetracycline (0.023 µM, Day 7), mercury chloride (3 µM, Day 8), vanillic acid (100 µM, Day 9), L-arabinose (6700 µM, Day 10), formaldehyde (25 µM, Day 11), and mercury chloride (3 µM, Day 12). Device effluent was collected daily via the integrated fraction collector, with a 400-µl sample taken for flow cytometry analysis. Samples were analyzed on the flow cytometer immediately following collection, and data were analyzed on FlowJo.

### Biosensor population distribution qPCR qPCR Primer Design

Primers for qPCR targeting each biosensor were designed using IDT primer quest and confirmed with BLAST to have less than 50% sequence match to BL21 genome. Following design, primers were tested for specificity to its corresponding biosensor by conducting qPCR with its biosensor in serial dilution of OD600. These data were used to generate calibration curves between CT value and log(OD600) of each biosensor. Primers were also tested with six other biosensors and BL21 negative control to confirm no unspecific primer binding. Each primer set was tested in a separate experiment with its corresponding MT. Primers that fit these requirements were deemed fit for upcoming population distribution experiments: negative controls that generated no CT value or CT value ≥34 and calibration curves with R2 ≥0.98.

### qPCR Sample Preparation and run

For tube culture population distribution experiments, cells were grown, combined, and passaged as mentioned in the “biosensor longevity test in batch culture” methods for the course of nine-day experiment. The co-cultures were induced on evenings on Days 1 (cuminic acid), 3 (L-arabinose), 6 (vanillic acid), and 8 (aTc). Samples (50 ul) were collected for qPCR analysis each day.

For qPCR sample preparation, cells were lysed by incubating at 95°C for 10 minutes, followed by storage at -20C before qPCR experiment. qPCR experiments probing for each biosensor were run separately to ensure optimal MT value used. Each qPCR experiment contained corresponding biosensor monoculture serial dilutions (for calibration curve generation), tested samples, and water control.

### Population distribution calculations

For population quantifications, OD600 for each biosensor was calculated using calibration curve relating CT values to log(O600). Relative cell density (RCD) was calculated by dividing OD600 for biosensor of interest by total OD600 of all biosensors (Equation 1).

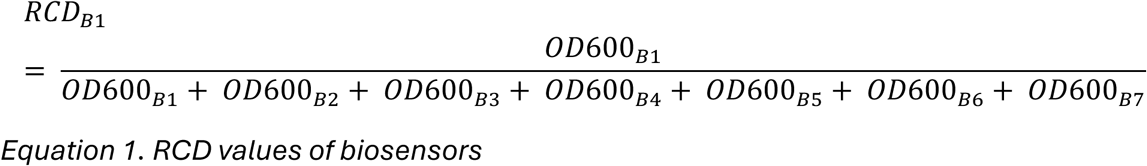

DCDs were calculated by subtracting day 1 RCD from consecutive day RCD values to demonstrate population variation over time.

### Vanilla extract dose-response validation in batch culture

Biosensor B3 was grown and induced as described above (“Cell culture and induction conditions”), using pure vanillic acid (100 µM) as the “Induced” positive control and no analyte as the “Uninduced” negative control. To assess response to a real-world sample, vanilla extract [365 by Whole Foods Market, Organic Vanilla Extract, 2 Fl oz] was added at increasing volumes (1, 5, 15, 17, 20, 40, and 100 µl) to separate wells in place of pure vanillic acid. Plates were read overnight on the plate reader (SpectraMax M5) as described above, and RFU/OD₆₀₀ was calculated for each condition. Statistical significance relative to the uninduced control was assessed by Welch’s t-test with Holm–Bonferroni correction for multiple comparisons (Fig. 5b). For fluorescence microscopy validation (Fig. 5c), B3 was induced overnight in batch monoculture with either pure vanillic acid or vanilla extract and imaged on the Echo Revolve microscope as described above.

### Multiplexed detection of vanilla extract in the mother machine

Starter cultures for each of the seven biosensors were grown, diluted, and combined at equal ratios as described in “Biosensor longevity test in mother machine” above, and loaded into the mother machine device at a combined OD₆₀₀ of 0.2. Defined medium was delivered by continuous flow at 3 µl/min throughout the two-day experiment. Pure vanillic acid (100 µM) was introduced into the flowing medium on Day 1 (morning), followed sequentially by increasing volumes of vanilla extract: 1 µl (Day 1, evening), 17 µl (Day 2, morning), and 100 µl (Day 2, evening). Device effluent was collected at each timepoint via the integrated fraction collector and analyzed by flow cytometry immediately following collection, using the gating and compensation workflow described in “Mother Machine data analysis for longevity experiment” above. Representative fluorescence micrographs (Fig. 5f) were acquired on the Echo Revolve microscope as described above.

## Acknowledgements

We thank Suckjoon Jun (UC San Diego) for generously providing PDMS replica. We thank Elizabeth Libby for generously providing plasmids encoding fluorescent proteins (YFP, CFP, mCherry) and ptrc99a backbone. We thank Aden Tom and Emma Gerlach for assisting with weekly QC of our WCBs and molecular cloning.

