## Supplemental Information for "Biosynthetic Olfactory System from Multiplexed Engineered Microbes (BOSMEM)"

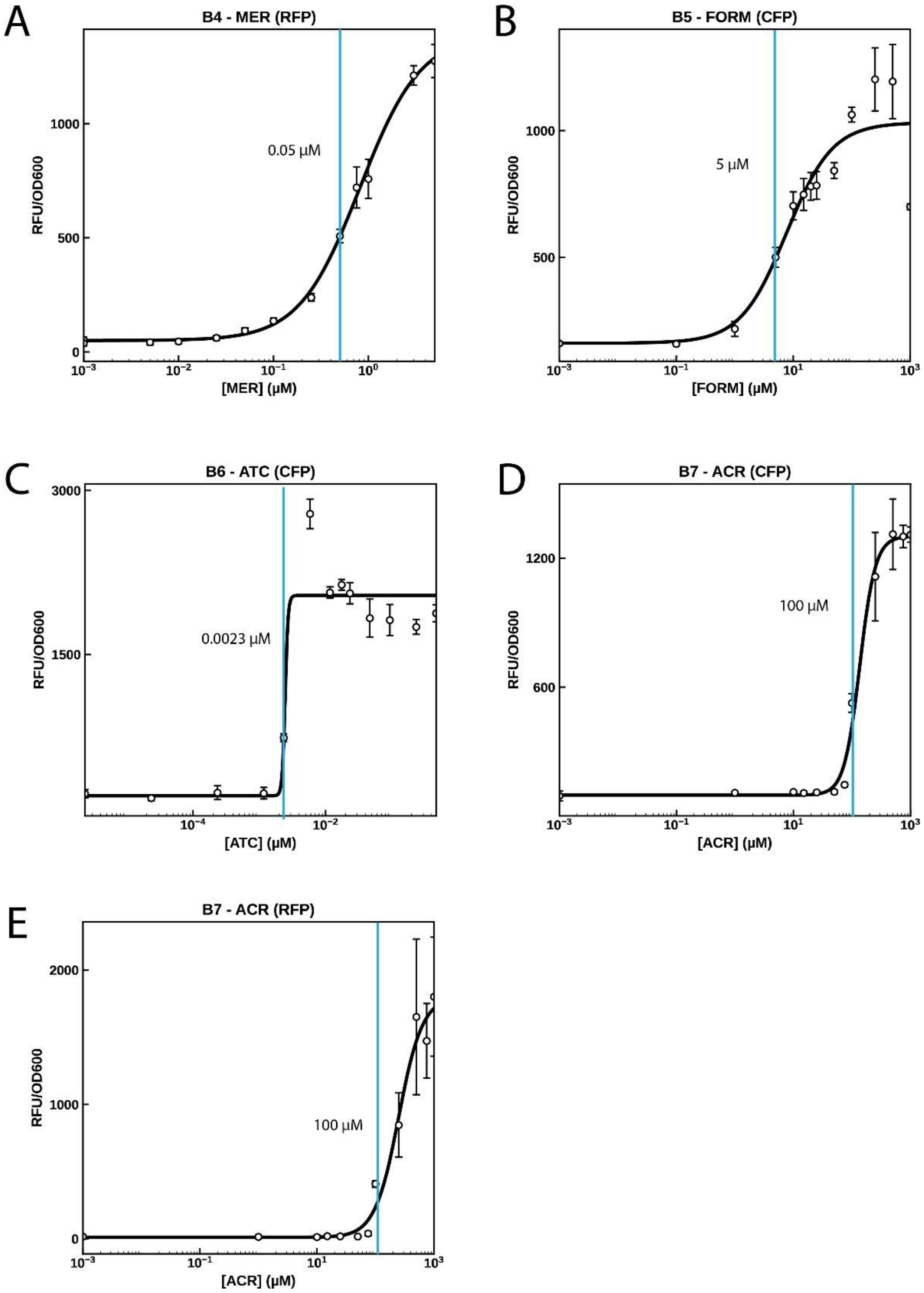


**Fig. S1 | Dose dependent curves for biosensors B4-7 in other FP channels.** Detection limit is noted by blue line.

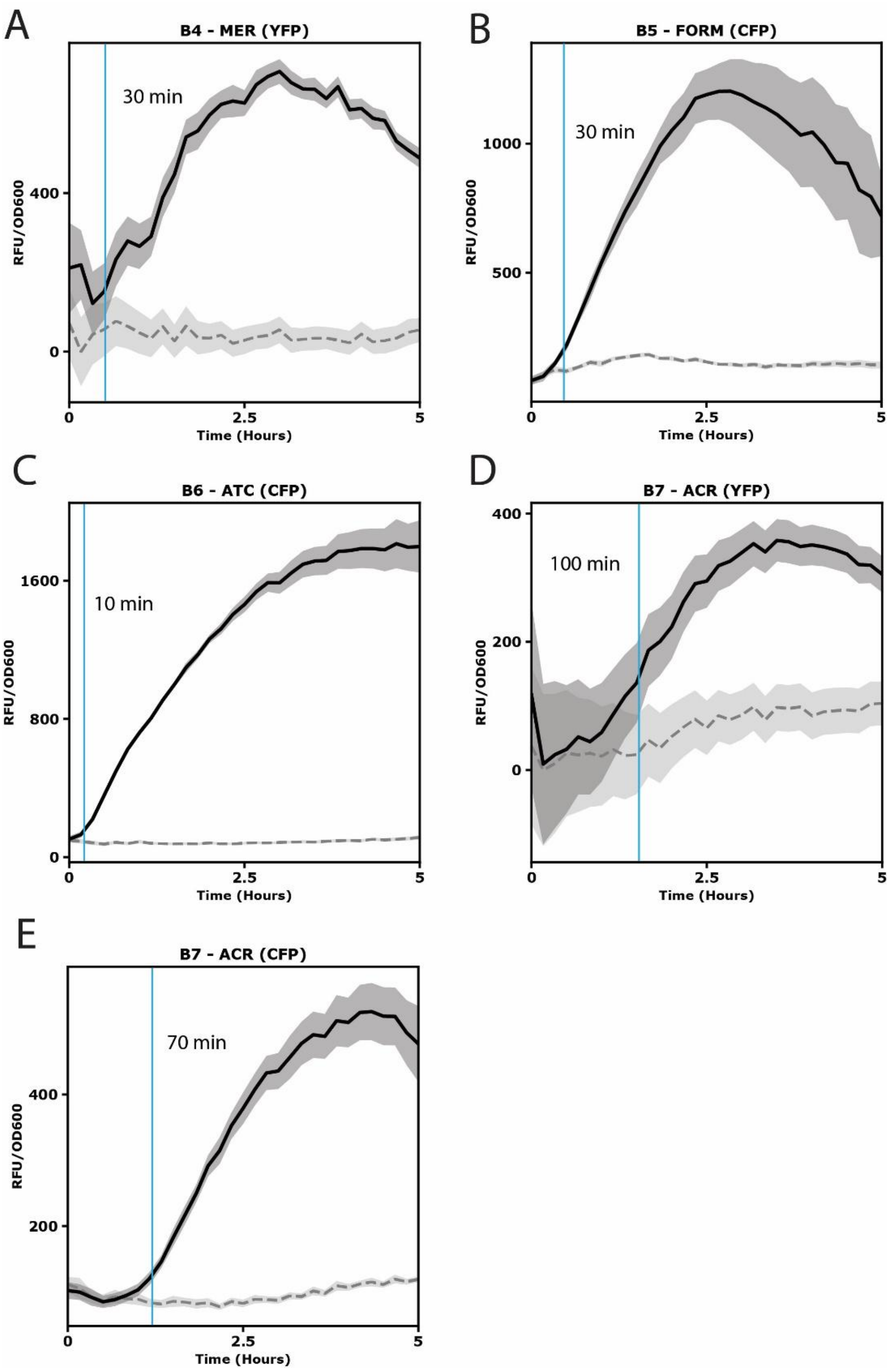


**Fig. S2 | Kinetic runs for biosensors B4-7 in other FP channels.** Activation time is noted by the blue line.

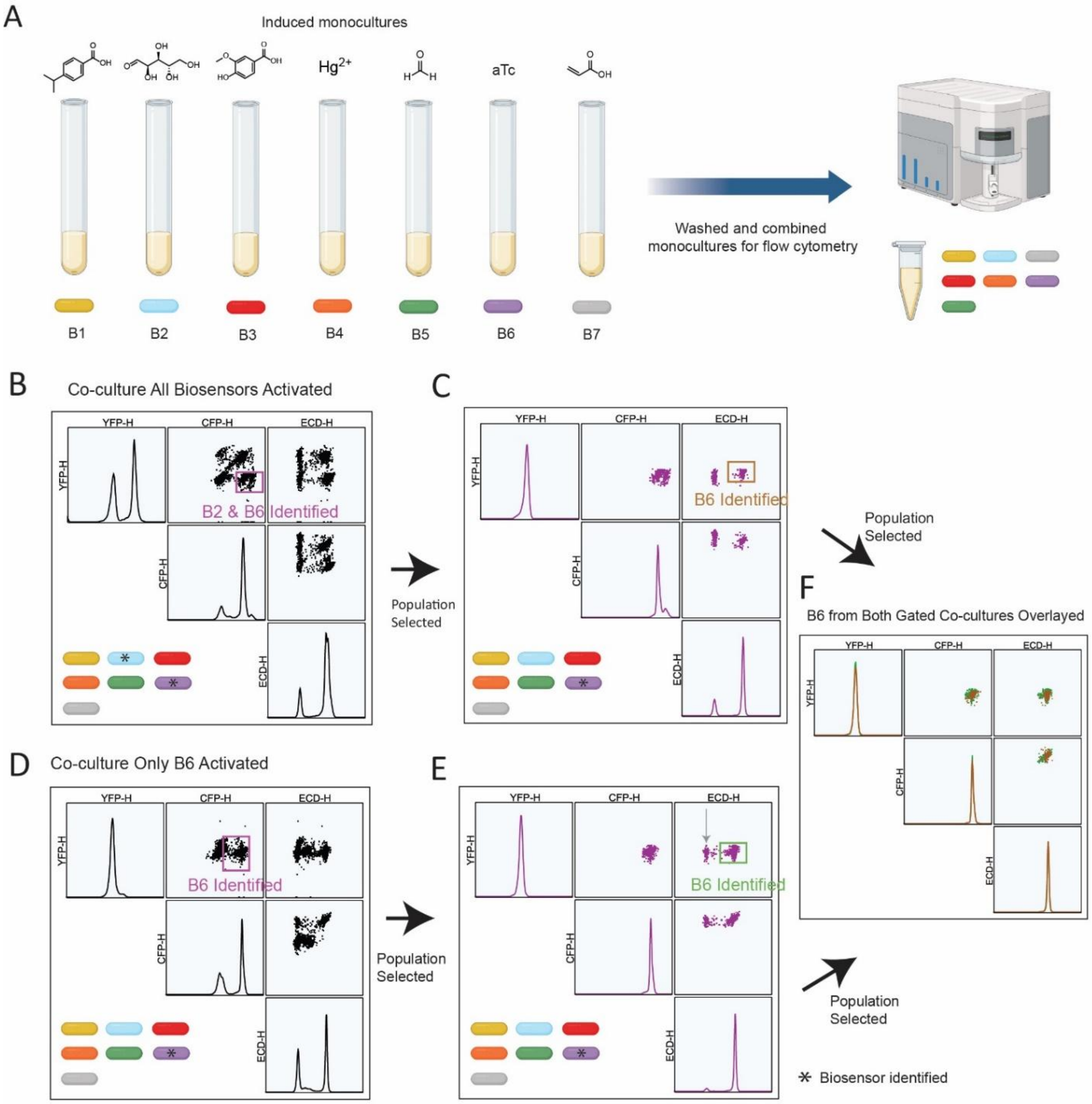


**Fig. S3 | Multiplex-ability of Biosensors in the Flow Cytometer. (A)** Sample preparation workflow for biosensors used in multiplex-ability analysis in flow cytometer. **(B-F)** Multiplex analysis to identify B6 population via flow cytometry gating. **(B)** Co-culture sample with all biosensors activated first gated by identifying a CFP+, YFP- population. Selected gate further analyzed by selecting RFP+ population **(C)**. Same analysis was performed on co-culture sample with only B6 activated **(D-E)**. To confirm B6 identification, B6 populations were overlayed **(F)**. Leaky expression of B2 in "Co-culture Only B6 Activated" sample signified by gray arrow.

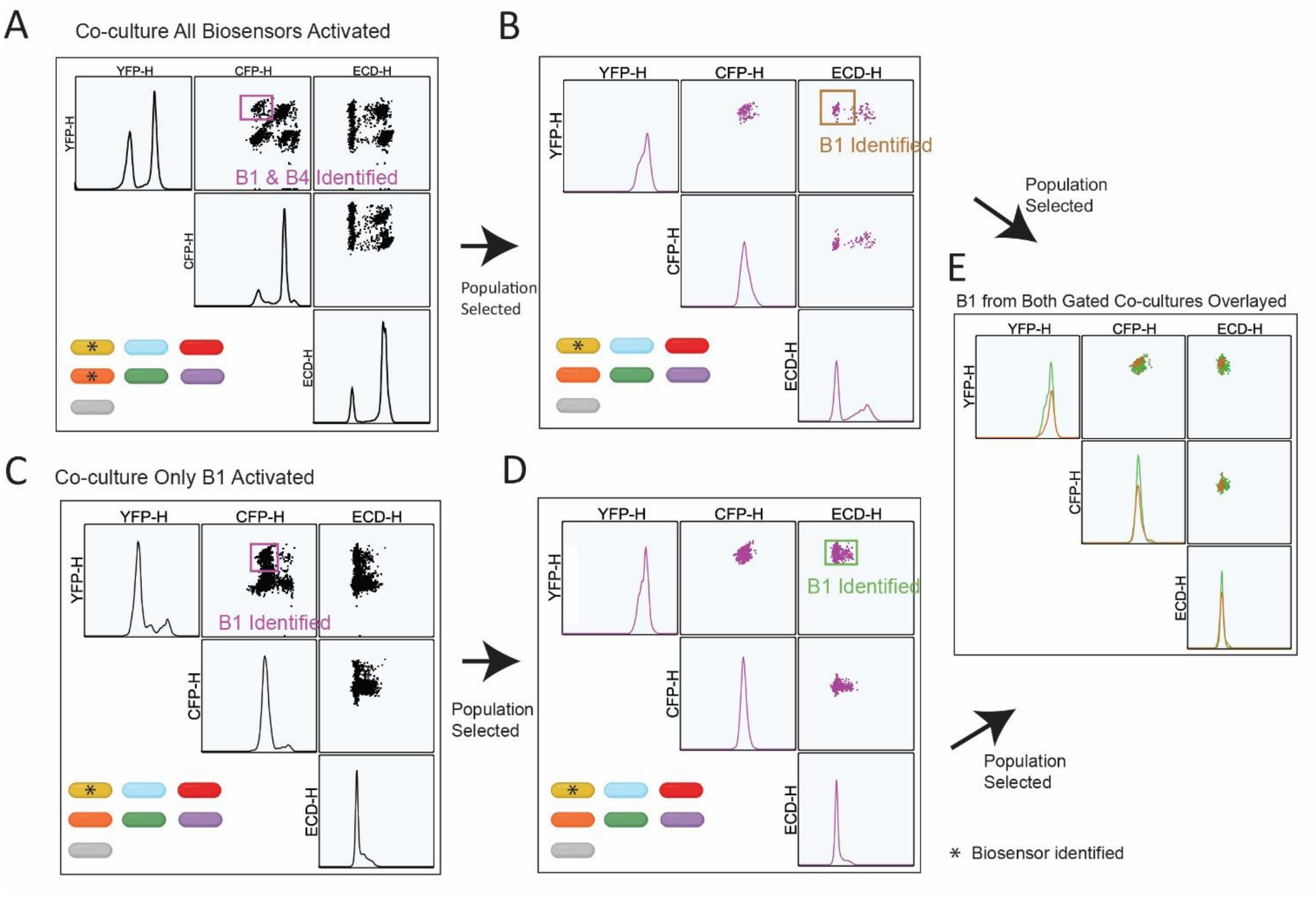


**Fig. S4. |** Multiplex analysis to identify B1 population via flow cytometry gating. **(A)** Co-culture sample with all biosensors activated first gated by identifying a YFP+, CFP- population. Selected gate further analyzed by selecting YFP+, RFP- population **(B)**. Same analysis was performed on co-culture sample with only B1 activated **(C-D)**. To confirm B1 identification, B1 populations were overlayed **(E)**.

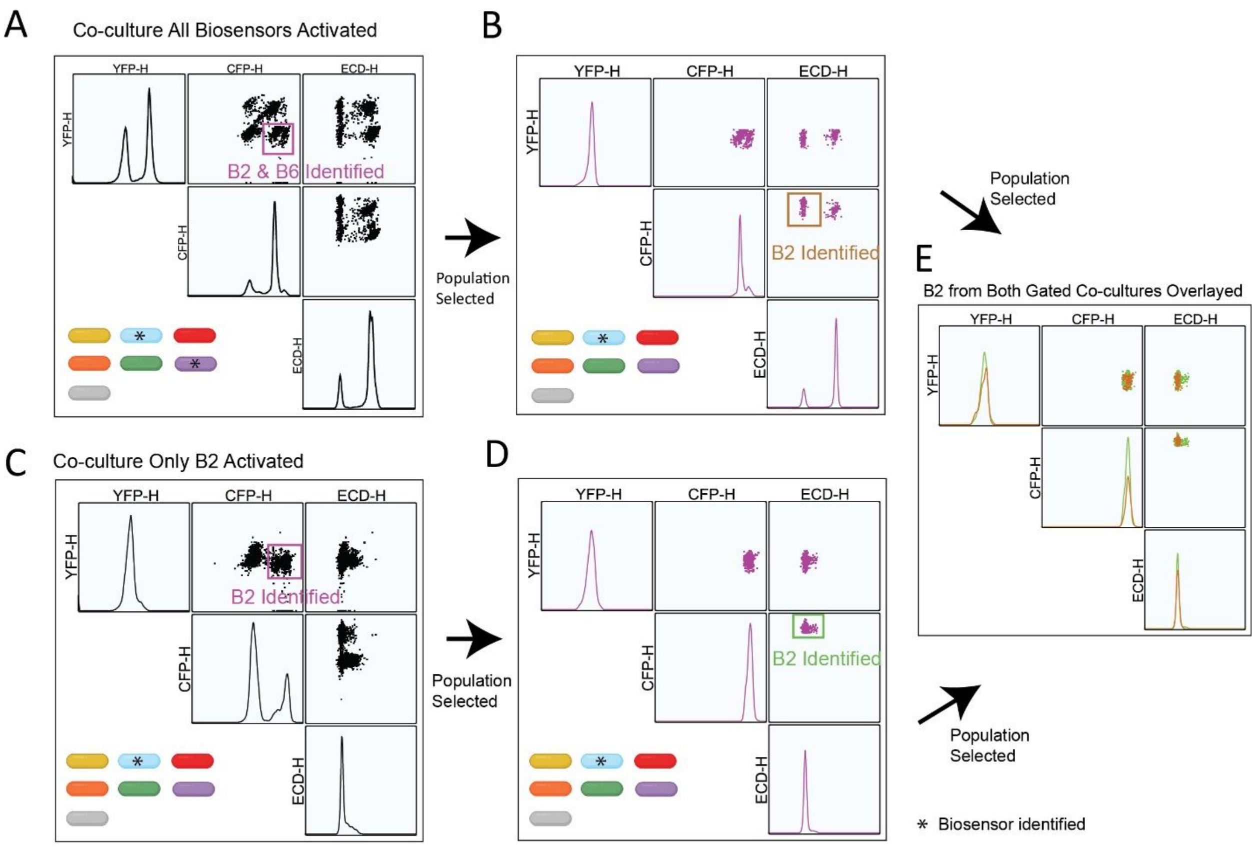


**Fig. S5. |** Multiplex analysis to identify B2 population via flow cytometry gating. **(A)** Co-culture sample with all biosensors activated first gated by identifying a CFP+, YFP- population. Selected gate further analyzed by selecting CFP+, RFP- population **(B)**. Same analysis was performed on co-culture sample with only B2 activated **(C-D)**. To confirm B2 identification, B2 populations were overlayed **(E)**.

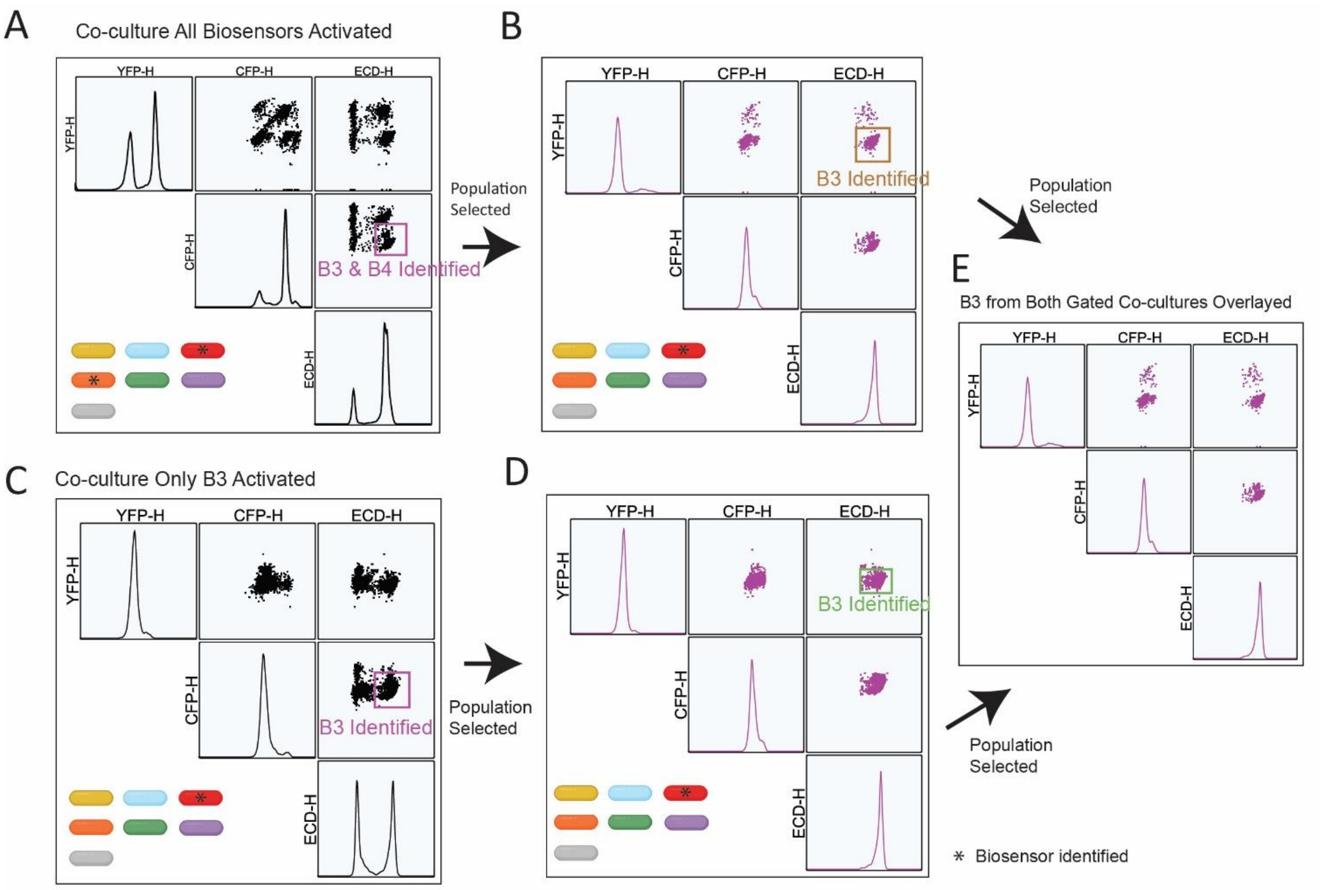


**Fig. S6. |** Multiplex analysis to identify B3 population via flow cytometry gating. **(A)** Co-culture sample with all biosensors activated first gated by identifying a RFP+, CFP- population. Selected gate further analyzed by selecting RFP+, YFP- population **(B)**. Same analysis was performed on co-culture sample with only B3 activated **(C-D)**. To confirm B3 identification, B3 populations were overlayed **(E)**.

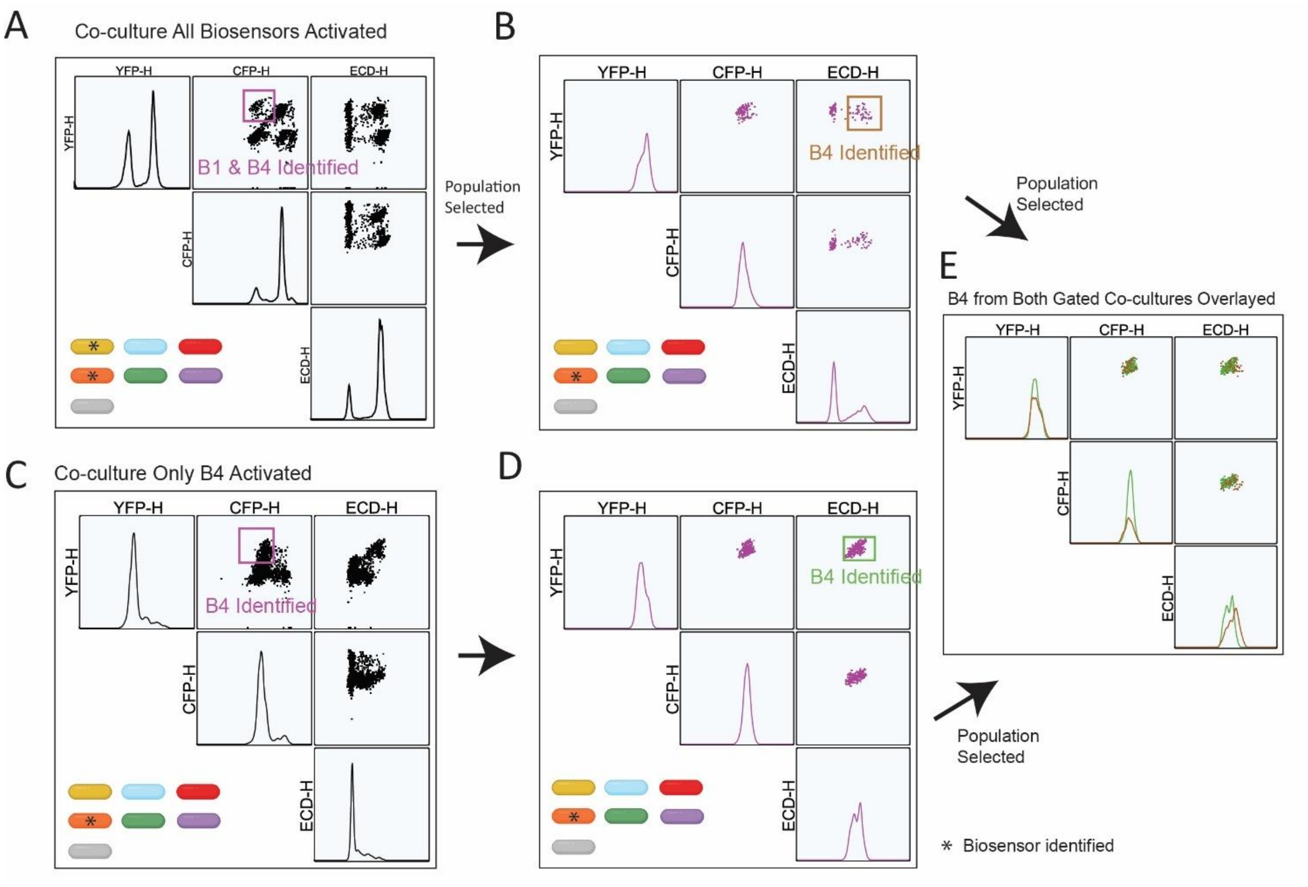


**Fig. S7. |** Multiplex analysis to identify B4 population via flow cytometry gating. **(A)** Co-culture sample with all biosensors activated first gated by identifying a YFP+, CFP- population. Selected gate further analyzed by selecting YFP+, RFP+ population **(B)**. Same analysis was performed on co-culture sample with only B4 activated **(C-D)**. To confirm B4 identification, B4 populations were overlayed **(E)**.

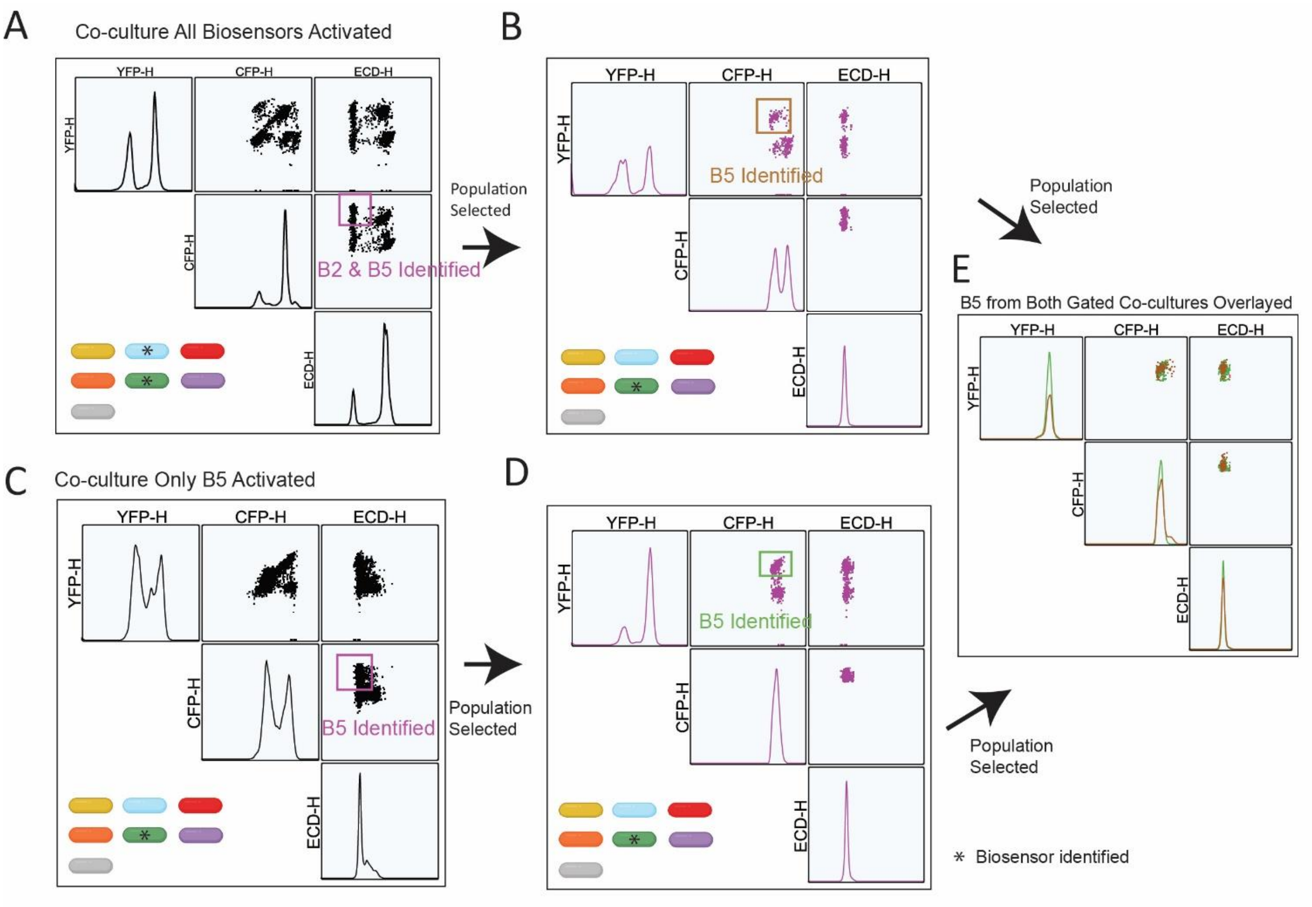


**Fig. S8. |** Multiplex analysis to identify B5 population via flow cytometry gating. **(A)** Co-culture sample with all biosensors activated first gated by identifying a CFP+, ECD- population. Selected gate further analyzed by selecting YFP+, CFP+ population **(B)**. Same analysis was performed on co-culture sample with only B5 activated **(C-D)**. To confirm B5 identification, B5 populations were overlayed **(E)**.

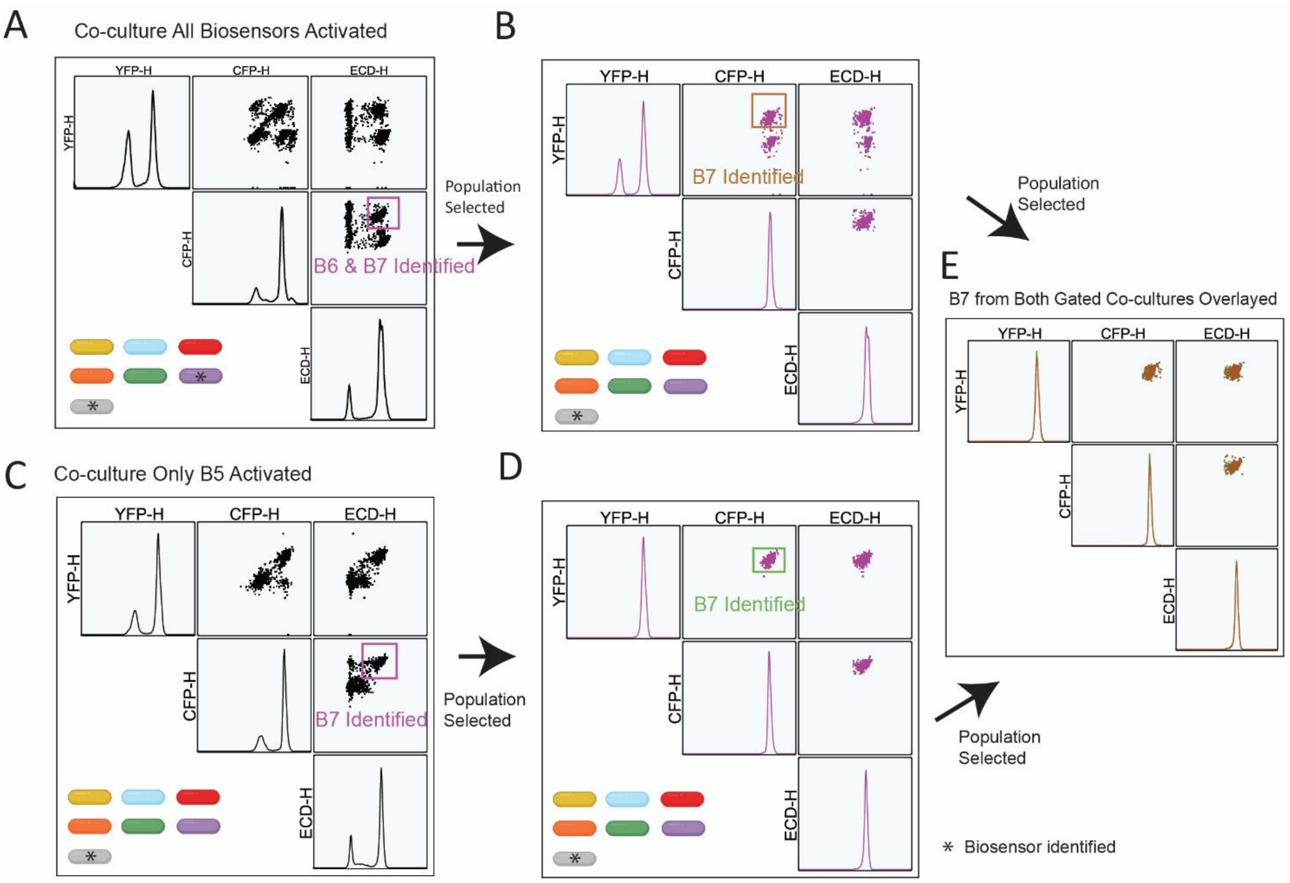


**Fig. S9. |** Multiplex analysis to identify B7 population via flow cytometry gating. **(A)** Co-culture sample with all biosensors activated first gated by identifying a CFP+, RFP+ population. Selected gate further analyzed by selecting YFP+, CFP+ population **(B)**. Same analysis was performed on co-culture sample with only B7 activated **(C-D)**. To confirm B7 identification, B7 populations were overlayed **(E)**.

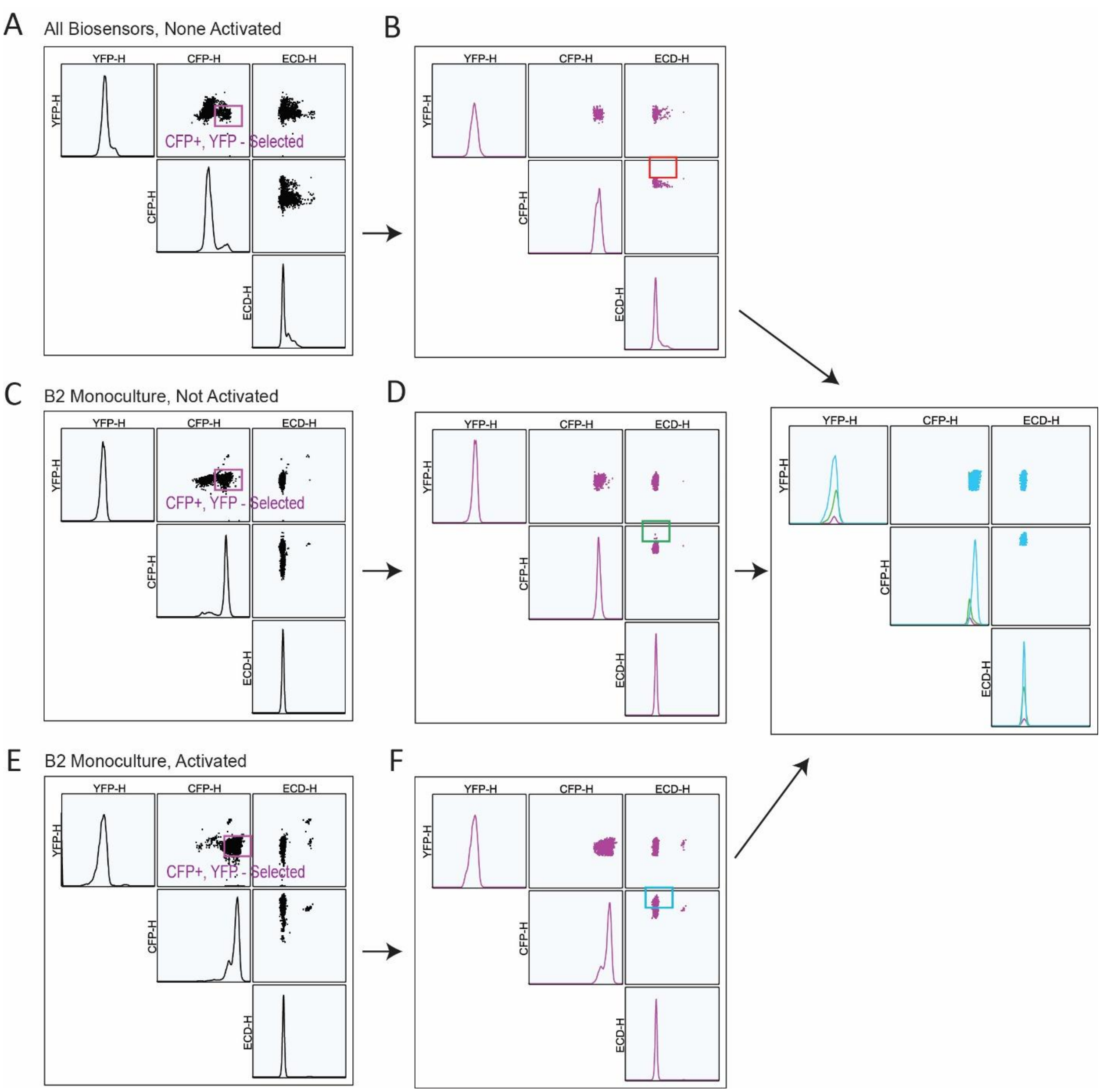


**Fig. S10 | Multiplex analysis to confirm leaky expression of B2.** In sample containing all biosensors, none activated, there should be no CFP+ expression. However, a CFP+ population is still visible in the sample. This population was separated by first selecting CFP+, YFP- sample **(A)**. The selected population displayed on another NxN plot (**B**) showed population located lower in CFP signal compared to CFP+, YFP-, RFP- gate that is usually selected for activated B2 population. The same results can be seen in a B2 inactive monoculture sample **(C)**, where there is a CFP+ population that is located below CFP+, YFP-, RFP- gate that is usually selected for activated B2 population **(D).** This is different from the B2 activated monoculture sample **(E)**, which contains a population in the B2 activated gate **(F)**. These results show that there was leaky expression of CFP+ from unactive B2, that can still be distinguished from activated B2 population through gating.

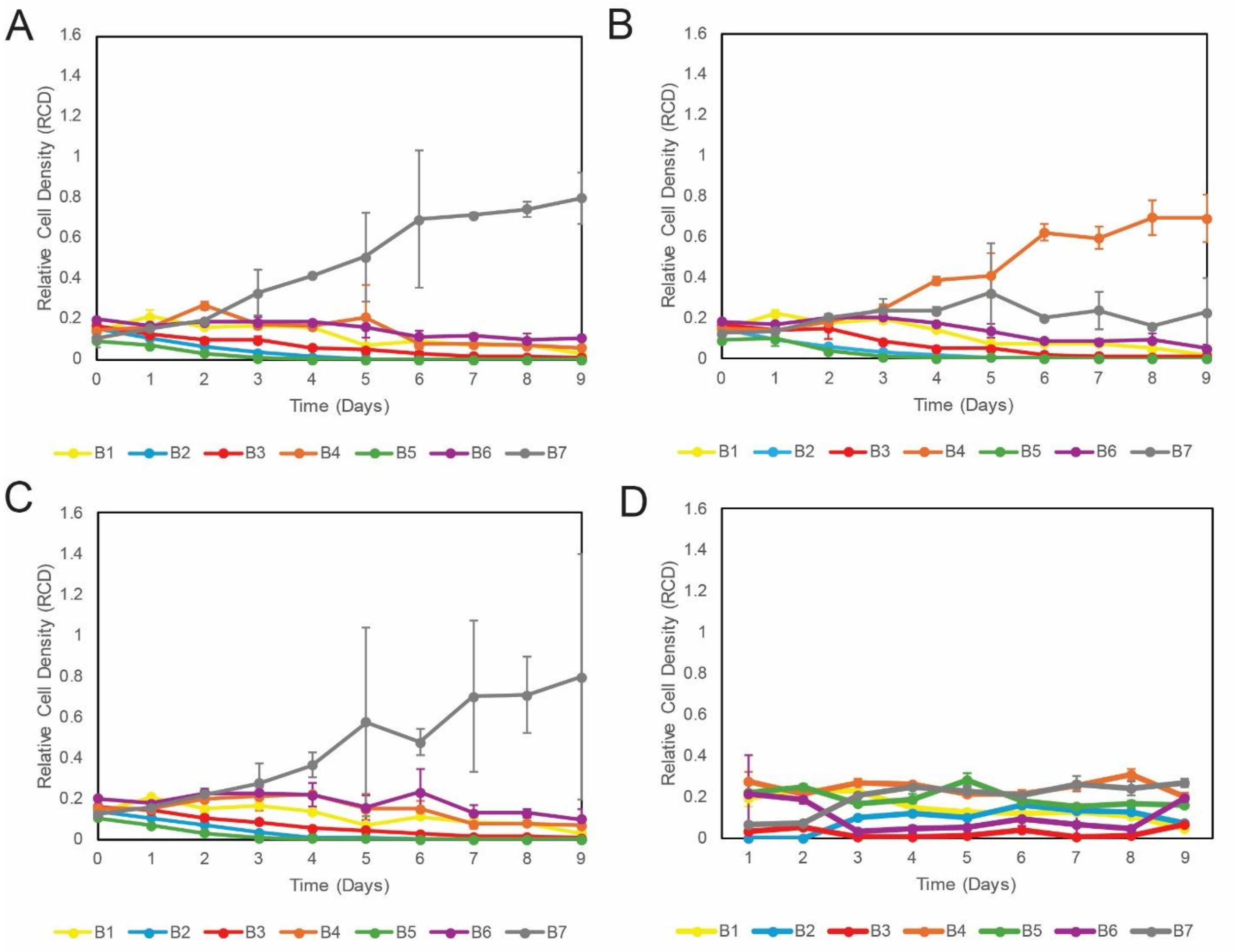


**Fig. S11. |** Relative cell densities (RCDs) for biosensors during longevity test for (A) original tube culture, (B) duplicate tube culture, (C) triplicate tube culture, and (D) mother machine effluent over the course of 9 days.

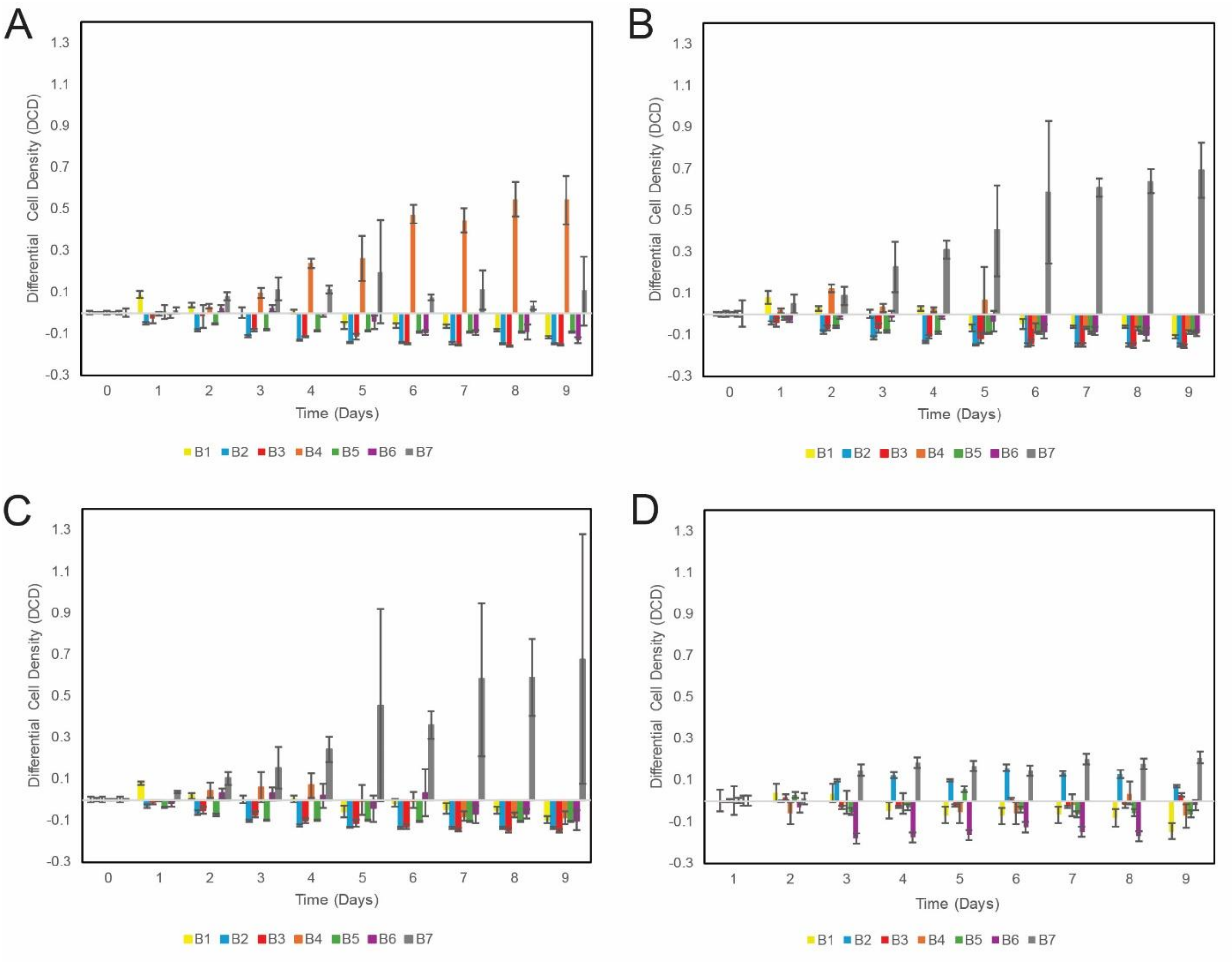


**Fig. S12. |** Differential cell densities (DCDs) for biosensors during longevity test for (A) original tube culture, (B) duplicate tube culture, (C) triplicate tube culture, and (D) mother machine effluent over the course of 9 days.

| Biosensor | Target Gene | Primer MT | Primers (5’ -> 3’) |
|---|---|---|---|
| B1 | cymR | 54 °C | Fwd: TCCAGCATCTGCTGAATAACATCATCT |
| | | | Rev: CTGGCAACCTTTGAATGGCTGTATG |
| B2 | tetA* | 56 °C | Fwd: CGCTGTAGGCATAGGCTTGGTTAT |
| | | | Rev: GACTGGCAATACTGTCGGAATGGA |
| B3 | vanR | 54 °C | Fwd: ACCAGGGTATCATGAAATGCCTGATTAT |
| | | | Rev: TGATGCAATTGAAGTTCGTGGTGTTC |
| B4 | merR | 57 °C | Fwd: GGCGCACACGAGTTCAGACA |
| | | | Rev: GGCCGAACACAAGCTCAAGGA |
| B5 | specR* | 56 °C | Fwd: CGGTGACCGTAAGGCTTGATGA |
| | | | Rev: GATGTCGTCGTGCACAACAATGG |
| B6 | tetR | 54 °C | Fwd: CGCCTTAGCCATTGAGATGTTAGATAGG |
| | | | Rev: TTTCTGTAGGCCGTGTACCTAAATGT |
| B7 | acuR | 55 °C | Fwd: GCTTCTCTTGCTCACCAGTCTCT |
| | | | Rev: GAACACGAGCAGCTTTCAGGATTTC |

**Table S1. | qPCR primer design and targeted genes.** *B2 and B5 use unique antibiotic resistance genes as target since repressor genes are also found in BL21 genome.

| | **CT Values Against Other Biosensors (Negative Control)** | | | | | | | |
|---|---|---|---|---|---|---|---|---|
| | **B1** | **B2** | **B3** | **B4** | **B5** | **B6** | **B7** | **BL21** |
| **B1** | -- | N/A | N/A | 38.30<br>37.20 | N/A | N/A | N/A | N/A |
| **B2** | N/A | -- | N/A | N/A | 35.56 | 36.47 | 34.69<br>37.11 | 35.51<br>40.41 |
| **B3** | N/A | N/A | -- | 36.57 | N/A | N/A | N/A | N/A |
| **B4** | N/A | N/A | N/A | -- | N/A | N/A | N/A | 40.82<br>40.80 |
| **B5** | N/A | N/A | N/A | N/A | -- | N/A | N/A | 34.63 |
| **B6** | N/A | N/A | N/A | N/A | N/A | -- | N/A | 34.63 |
| **B7** | 35.69 | 36.06 | 35.40 | N/A | 39.31 | N/A | -- | N/A |

**Table S2. | CT values testing biosensor primers against other biosensor negative controls.** Negative controls were tested at OD=0.0005.

| | Median Fold Change | | | Uninduced rCV | | | Induced rCV | | |
|---|---|---|---|---|---|---|---|---|---|
| | YFP | CFP | RFP | YFP | CFP | RFP | YFP | CFP | RFP |
| B1 | 87.9× | 1.1× | 1.0× | 76.5 | 79.6 | 23.6 | 58.3 | 44.9 | 23.0 |
| B2 | *0.8×* ▼ | 6.4× | 1.0× | 54.4 | 72.4 | 23.2 | 96.0 | 70.6 | 22.7 |
| B3 | 1.2× | 1.3× | 44.4× | 41.8 | 61.0 | 63.7 | 49.7 | 58.3 | 56.8 |
| B4 | 34.9× | 2.1× | 27.2× | 48.7 | 66.4 | 82.2 | 152.0 | 52.5 | 93.6 |
| B5 | 30.7× | 15.7× | 1.0× | 43.2 | 55.5 | 22.1 | 90.9 | 94.1 | 24.0 |
| B6 | 1.6× | 51.8× | 192.2× | 39.9 | 59.0 | 49.8 | 35.8 | 23.8 | 21.1 |
| B7 | 6.5× | 31.6× | 262.9× | 56.4 | 76.8 | 42.2 | 39.6 | 27.6 | 34.6 |

**Table S3. Robust coefficient of variation (rCV) and median fold change in reporter expression across the biosensor library.**

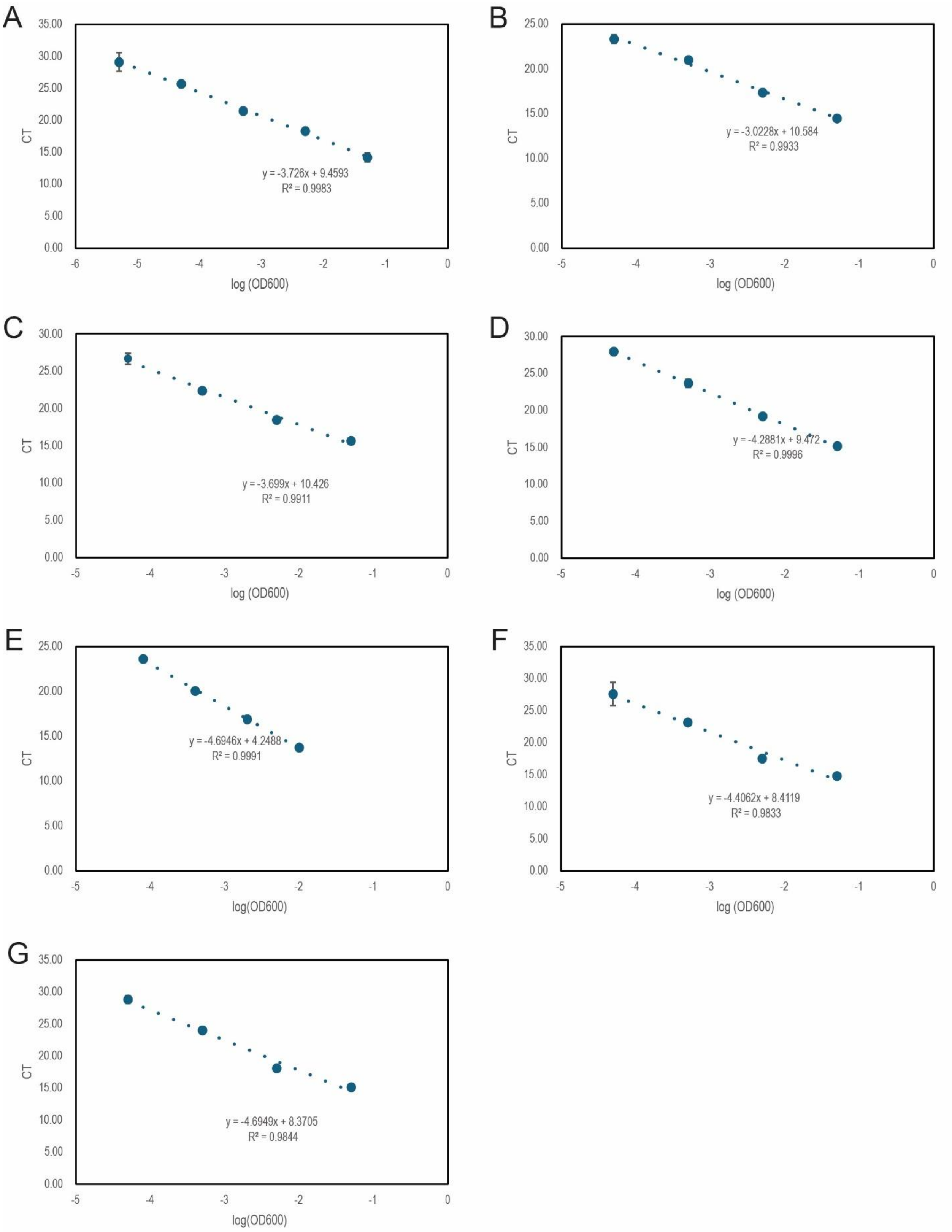


**Fig. S13. |** Calibration curves relating OD600 values of biosensors (A) B1, (B) B2, (C) B3, (D) B4, (E) B5, (F) B6, and (G) B7, to CT values in qPCR.
